# PPARdelta signaling activation improves metabolic and contractile maturation of human pluripotent stem cell-derived cardiomyocytes

**DOI:** 10.1101/2021.07.12.451352

**Authors:** Nadeera M. Wickramasinghe, David Sachs, Bhavana Shewale, David M. Gonzalez, Priyanka Dhanan-Krishnan, Denis Torre, Elizabeth LaMarca, Serena Raimo, Rafael Dariolli, Madhavika N. Serasinghe, Joshua Mayourian, Robert Sebra, Kristin Beaumont, Ravi Iyengar, Deborah L. French, Arne Hansen, Thomas Eschenhagen, Jerry E. Chipuk, Eric A. Sobie, Adam Jacobs, Schahram Akbarian, Harry Ischiropoulos, Avi Ma’ayan, Sander M. Houten, Kevin Costa, Nicole C. Dubois

## Abstract

Pluripotent stem cell-derived cardiomyocytes (PSC-CMs) provide an unprecedented opportunity to study human heart development and disease. A major caveat however is that they remain functionally and structurally immature in culture, limiting their potential for disease modeling and regenerative approaches. Here, we address the question of how different metabolic pathways can be modulated in order to induce efficient hPSC-CM maturation. We show that PPAR signaling acts in an isoform-specific manner to balance glycolysis and fatty acid oxidation (FAO). PPARD activation or inhibition results in efficient respective up- or down-regulation of the gene regulatory networks underlying FAO in hPSC-CMs. PPARD induction further increases mitochondrial and peroxisome content, enhances mitochondrial cristae formation and augments FAO flux. Lastly PPARD activation results in enhanced myofibril organization and improved contractility. Transient lactate exposure, commonly used in hPSC-CM purification protocols, induces an independent program of cardiac maturation, but when combined with PPARD activation equally results in a metabolic switch to FAO. In summary, we identify multiple axes of metabolic modifications of hPSC-CMs and a role for PPARD signaling in inducing the metabolic switch to FAO in hPSC-CMs. Our findings provide new and easily implemented opportunities to generate mature hPSC-CMs for disease modeling and regenerative therapy.

## INTRODUCTION

The differentiation of human pluripotent stem cells (hPSCs) to various cell types of the human body has opened unprecedented opportunities to study the biology of human cells and tissues *in vitro*. This is particularly relevant for studying the heart, as heart disease continues to be the leading cause of death worldwide but access to tissue samples and establishment of relevant animal models remains challenging (Brandão et al., 2017; Hasenfuss, 1998; Houser et al., 2012; Tsang et al., 2016; Virani et al., 2020). Cardiomyocytes (CMs) can be derived with high efficiency from hPSCs, and have been used successfully for disease modeling, regenerative therapies and drug discovery (Burridge et al., 2015; Kattman et al., 2011; Lian et al., 2012, 2013; Murry and Keller, 2008; Yang et al., 2008; Zhang et al., 2012). However, studies assessing cell morphology, gene expression, electrophysiological properties, mitochondrial content, metabolic activity, contractile force and formation of T-tubules demonstrate that hPSC-CMs remain functionally and structurally immature, limiting their therapeutic and research use (van den Berg et al., 2015; DeLaughter et al., 2016; Gherghiceanu et al., 2011; Karbassi et al., 2020; Liu et al., 2010; Parikh et al., 2017; Thavandiran et al., 2013; Ulmer and Eschenhagen, 2020; Yang et al., 2014a). Current differentiation protocols to generate hPSC-CMs are largely based on recapitulating *in vitro* the developmental stages of *in vivo* heart development by providing the appropriate signaling environments (Calderon et al., 2016; Kattman et al., 2011; Lian et al., 2012, 2013; Yang et al., 2008). Thus hPSC-CM immaturity is likely the result of an incomplete recapitulation of heart development in current *in vitro* PSC differentiation protocols.

Cardiac maturation after midgestation is dependent on the coordinated development of metabolic, electrochemical, mechanical and cell-cell interaction mechanisms. Consequently, strategies to enhance hPSC-CM maturation have focused on long-term cultures (DeLaughter et al., 2016; Kamakura et al., 2013), tissue engineering (Correia et al., 2018; Giacomelli et al., 2020; Leonard et al., 2018; Li et al., 2018; Lopez et al., 2021; Nunes et al., 2013; Shadrin et al., 2017; Thavandiran et al., 2013; Tiburcy et al., 2017; Zhao et al., 2020), electrical stimulation (Nunes et al., 2013; Ronaldson-Bouchard et al., 2018), activation of developmentally-relevant signaling pathways (Gentillon et al., 2019; Kosmidis et al., 2015; Lopez et al., 2021; Murphy et al., 2021; Parikh et al., 2017; Poon et al., 2015; Yang et al., 2014b), epigenetic modifications and substrate utilization (Correia et al., 2017; Feyen et al., 2020; Hu et al., 2018; Mills et al., 2017; Nakano et al., 2017; Yang et al., 2019). Several of these approaches were successful in enhancing hPSC-CM maturation, yet the mechanisms impeding maturation beyond fetal stages are still not fully understood and are likely to be driven by a combination of new and aforementioned parameters.

In addition to the numerous complex morphological and structural changes that occur as the heart grows, the embryo also encounters systemic changes, one of them being the well-characterized metabolic switch (Ascuitto and Ross-Ascuitto, 1996; Lopaschuk and Jaswal, 2010; Piquereau and Ventura-Clapier, 2018). The adult heart predominantly utilizes mitochondrial fatty acid oxidation (FAO) to fuel ATP production (Houten et al., 2016; Stanley et al., 2005). In contrast, the developing heart utilizes anaerobic glycolysis and lactate oxidation to generate ATP (Lopaschuk and Jaswal, 2010; Piquereau and Ventura-Clapier, 2018). Although a marked metabolic shift toward FAO occurs postnatally, upon the availability of dietary long chain fatty acids (LCFAs), the transcriptional program required for FAO is established before that, as early as mid-gestation (Menendez-Montes et al., 2016; Uosaki et al., 2015). Metabolic activity at different stages of development occurs in response to substrate and oxygen availability and changes in workload, with the first two being dependent on the expression of the machinery for glycolysis, glucose and lactate oxidation or FAO respectively. The metabolic switch allows the organism to become more flexible with respect to substrate use, and to meet the higher energy demands of the growing heart as FAO generates more ATP per mole of substrate oxidized than any other pathway (Lopaschuk and Jaswal, 2010; Piquereau and Ventura-Clapier, 2018).

The mechanisms that induce this metabolic switch during development are poorly understood. Recent in depth transcriptional profiling of the developing heart provide comprehensive resources to investigate the changes that occur as the heart matures (DeLaughter et al., 2016; Uosaki et al., 2015). Starting at midgestation the heart begins to express genes involved in LCFA metabolism and expression of FAO pathway genes continuously increases as development proceeds. The Peroxisome-Proliferator-Associated-Receptor (PPAR) pathway parallels this expression pattern and turns on at midgestation, increases continuously to then remain active throughout adulthood. PPARs belong to a class of ligand-activated transcription factors, nuclear hormone receptors, that are involved in growth, proliferation and metabolism. They exist in three isoforms, PPARalpha (PPARA), PPARbeta/delta (PPARD) and PPARgamma (PPARG) (Ahmadian et al., 2013; Barger and Kelly, 2000; Houten et al., 2016; Lopaschuk and Jaswal, 2010). Knockout (KO) mouse models for PPARA and PPARD have underscored their role in metabolic regulation. PPARA-deficient mice show reduced FAO gene expression, but fasting is required for this phenotype to become apparent (Liu et al., 2011). Deletion of PPARD results in embryonic lethality, however CM-restricted PPARD KO mice are viable with reduced FAO and increased glucose uptake in adulthood (Barak et al., 2002; Cheng et al., 2004). Based on these studies, we hypothesized that inducing PPAR signaling *in vitro* will model the contributions of the signaling pathway to the metabolic switch that is observed during *in vivo* heart development.

The overall goal of this study was to recapitulate the key metabolic changes that occur during *in vivo* heart development by modulation of substrate availability and activation of PPAR signaling. We uncovered that activation of PPAR signaling acts in an isoform-specific manner on hPSC-CMs to recapitulate key changes that occur during *in vivo* heart development. Activation of PPARD signaling specifically induces the expression of genes required for FAO across multiple culture formats and does so via sequential activation of gene regulatory networks over time of differentiation. These changes further result in increased peroxisomal and mitochondrial mass, enhanced cristae structure, and concordantly improve the ability of cells to utilize LCFAs as an energy substrate. PPARD signaling activation enhances mature cell morphology and myofibril alignment, and the number of bi-nucleated hPSC-CMs. Accordingly, increased PPARD activation leads to a prolonged action potential and increased contractility, both features of functional hPSC-CM maturation. Lastly, transient lactate exposure, frequently used to enrich for hPSC-CMs, induces an alternative program of cardiac maturation by activating components of cardiac contractility. The combination of lactate exposure and PPARD activation maintained both the metabolic and contractile transcriptional maturation phenotype. In conclusion, we identify an isoform-specific role for PPARD signaling in regulating the metabolic switch to FAO during early human heart development. Lactate selection further induces a parallel maturation program, jointly resulting in hPSC-CMs resembling a more mature stage of *in vivo* development. Together, we provide strategies to further refine existing differentiation protocols, in an easy and scalable manner, to ultimately yield physiologically relevant cardiac cells for *in vitro* studies and therapeutics development.

## RESULTS

### PPARD signaling enhances myofibril organization in hPSC-CMs in an isoform-specific manner

To first assess PPAR signaling activity during hPSC differentiation we performed qPCR analysis for the three PPAR isoforms, PPARA, PPARD and PPARG. Gene expression of all isoforms continuously increased over the course of differentiation, but remained low compared to 20-week-old fetal and adult heart tissue, suggesting that current differentiation protocols may not recapitulate the PPAR signaling activity observed in the human heart (**Figure 1A**). Immunofluorescence analysis of 16-week-old human fetal heart confirmed the gene expression analysis and reflects the typical PPAR isoform distribution described in the developing mouse heart (**Figure 1B** and **Figure S1A**) (Uosaki et al., 2015). To explore a potential requirement for PPAR signaling during cardiac differentiation we inhibited PPAR signaling in hPSC-CMs using highly selective small molecule antagonists for each of the three isoforms (PPARA: GW6471, PPARD: GSK0660 (Shearer et al., 2008), PPARG: GW9662). Abrogation of PPAR signaling throughout differentiation, individually or in combination, did not affect the generation of hPSC-CMs, as measured by the percentage of SIRPA+CD90-hPSC-CMs at day 20 of differentiation (**Figure 1C/F)**. However, the combination of all isoform-specific antagonists reduced the number of total cells, suggesting that PPAR signaling is not required for CM specification, but may play a role in cell expansion during early development (**Figure 1G**). Next, we tested PPAR inhibition during later stages of CM differentiation, when PPAR expression levels are increased. hPSC-CMs were plated as 2D monolayers at day 20 of differentiation, followed by two weeks of PPAR antagonist treatment. Flow cytometry analysis for SIRPA+CD90+ cells showed no difference in hPSC-CM maintenance after PPAR inhibition (**Figure 1D/H**). However, analysis of alpha-actinin (ACTN2) expression using an ACTN2-eGFP PSC reporter line revealed that inhibition of all three PPAR isoforms results in decreased ACTN2 expression in hPSC-CMs (**Figure 1E/I/J**). IF analysis for the sarcomere proteins ACTN2 and cardiac Troponin T (cTNT) further illustrated that inhibition of PPARA and of all PPAR isoforms combined has adverse effects on hPSC-CM morphology with fewer and less organized myofibrils (**Figure 1K**). Inhibition of PPARD or PPARG did not impact hPSC-CM morphology. To understand whether these isoform-specific effects correlate with overall abundance of individual PPARs we performed single cell RNA sequencing (scRNAseq) of hPSC-CMs and assessed the expression of all three PPAR isoforms. As expected, we found PPARs to be expressed primarily in hPSC-CMs and not in fibroblast cells (**Figure 1L/M**). PPARA was the most abundantly expressed isoform, followed by PPARD and low levels of PPARG expression (**Figure 1M/N**). This pattern was further confirmed by IF analysis in hPSC-CMs (**Figure S1C**). Lastly, we confirmed expression of PPAR signaling co-activators (PPARGC1A, PPARGC11B) and the PPAR heterodimer binding partners Retinoid X Receptors (RXRA, RXRB and RXRC) in hPSC-CMs, collectively suggesting that the PPAR signaling machinery is present in hPSC-CMs, and that PPARA is the predominant isoform in hPSC-CMs (**Figure S1D-F**).

**Figure 1:**
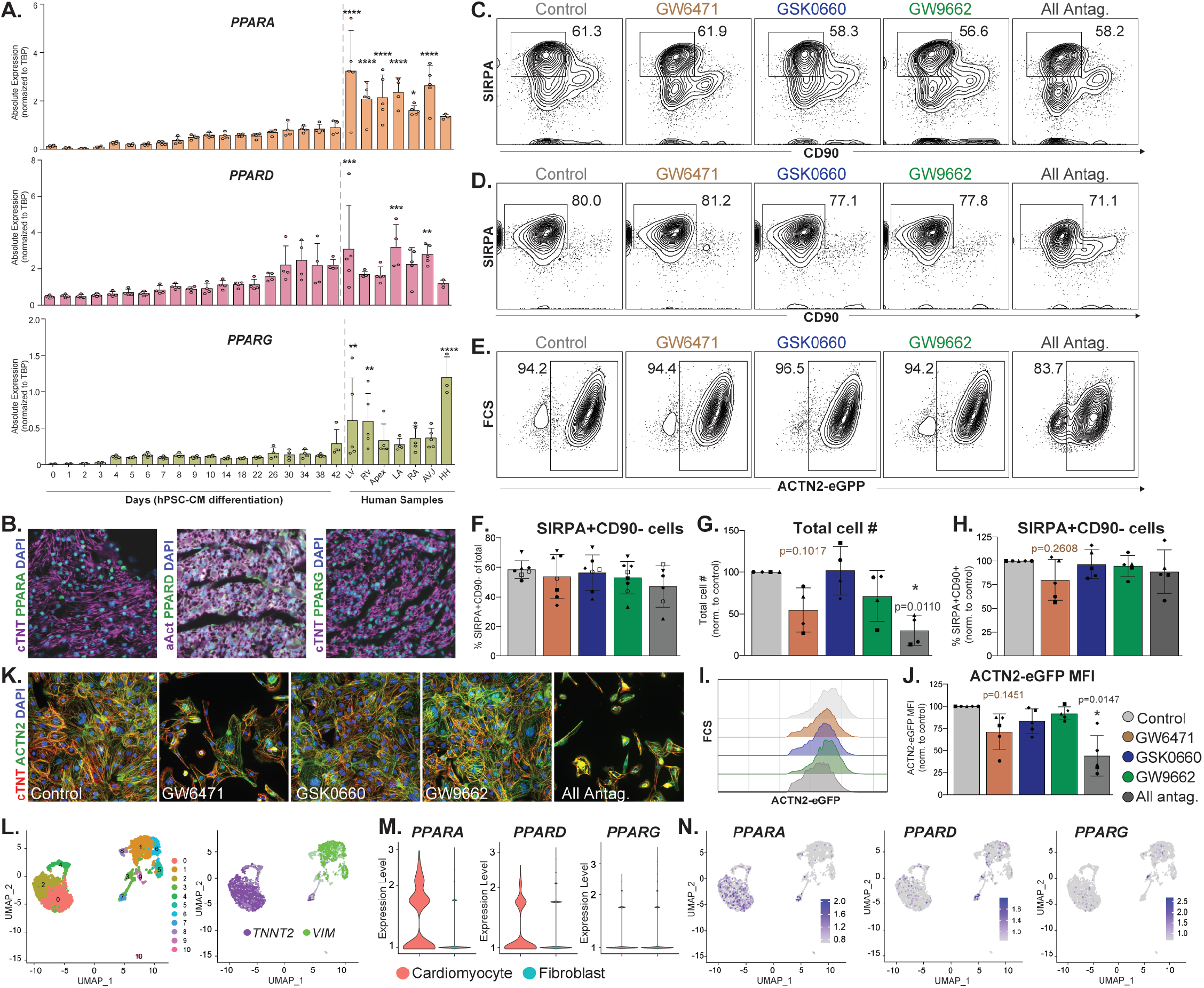
PPAR isoforms show distinct expression and function during hPSC-CM differentiation and maturation. (**A**) RT-qPCR analysis of *PPARA, PPARD* and *PPARG* during hPSC-CM differentiation (n=4) and in human fetal heart tissue (week 20 of gestation). (**B**) IF analysis for cardiac Troponin T (cTnT) or alpha-Actinin (aAct) in combination with PPARA, PPARD or PPARG on week 16 human fetal heart (left ventricle). DAPI is used to visualize nuclei. Scale bar = 25μm. (**C**) Flow cytometry analysis of hPSC-CMs at day 20 of differentiation for SIRPA (cardiomyocytes) and CD90 (fibroblast cells). (**D/E**) Flow cytometry analysis of hPSC-CMs at day 35 of differentiation (20 days as EBs, followed by 15 days in 2D) for SIRPA (cardiomyocytes) and CD90 (fibroblast cells), or for ACTN2-eGFP. Cells were treated with PPAR antagonists for 2 weeks prior to analysis. (**F**) Quantification of SIRPA+CD90-hPSC-CMs from (**C**). (**G**) Quantification of total cell numbers from (C). (**H**) Quantification of SIRPA+CD90-hPSC-CMs from (D). (**I**) ACTN2-eGFP fluorescence intensity in hPSC-CMs treated with PPAR antagonists for 2 week (days 20-35). (J) Quantification of ACTN2-eGFP mean fluorescence intensity in (I). (**K**) IF analysis for cardiac Troponin T (cTnT) and alpha-Actinin (aAct) in hPSC-CMs treated with PPAR antagonists for 2 week (days 20-35). DAPI is used to visualize nuclei. Scale bar = 25μm. (**L**) ScRNAseq UMAP clustering of untreated hPSC-CMs (left) and expression of cTnT (green) and Vimentin (purple)(right). (**M**) cTnT+ (red) and Vimentin+ (blue) cells were grouped, and expression of PPAR isoforms assessed in both populations. (**N**) Expression of *PPARA, PPARD* and *PPARG* in hPSC-CMs. Biological replicates are identified by same shape of respective data points. Data represented as mean + SD. Statistics: Student’s t-tests relative to the control condition (far left) (*p<.05; **p<.01; ***p<.001; **** p<.0001). AVJ: Atrioventricular junction, HH: Human heart (adult), LA: left atria, LV: left ventricle, RA: right atrial, RV: right ventricle.

To test the effect of PPAR signaling activation we treated differentiated hPSC-CMs with isoform-specific small molecule agonists (PPARA: WY14643, PPARD: GW0742 (Sznaidman et al., 2003), PPARG: Rosiglitazone) for 4 weeks and assessed changes in cell morphology and sarcomere organization (**Figure 2A**). Immunofluorescence (IF) analysis of cardiac Troponin T (cTnT) revealed an isoform-specific effect on myofibril alignment in hPSC-CMs treated with GW0742 (PPARD), but not WY14643 (PPARA) or Rosiglitazone (PPARG)(Figure 2B). To quantify myofibril organization in an unbiased manner images of cTnT-stained hPSC-CMs were processed using MatFiber, a vector-based program (**Figure S2A**) (Fomovsky and Holmes, 2010). This analysis revealed that myofibrils of PPARD-activated hPSC-CMs, but not PPARA or PPARG, exhibit greater alignment against each other and along the length of the cell, together with a reduced circularity standard deviation, both of which are characteristic of enhanced CM structural maturation (**Figure 2C/D**). However, PPARD activation did not affect sarcomere lengths or regularity, as assessed by the distance between Z-discs labeled by alpha actinin (**Figure S2B/C**). The developing mammalian heart progresses through a phase of extensive proliferation before birth, followed by hypertrophic growth and a reduction in the proliferative capacity of mature CMs. To assess the effect of PPAR signaling on hPSC-CM proliferation, we performed EdU incorporation assays after 4 weeks of PPAR activation/inhibition. The percentage of EdU positive cells was unchanged in all culture conditions after continuous PPAR manipulation, indicating that proliferation capacity is not altered by activation/inhibition of PPAR signaling in hPSC-CMs (**Figure S2D**). To assess hPSC-CM cell size we measured the total surface area of alpha actinin-positive hPSC-CMs 4 weeks after PPAR activation or inhibition. Consistent with increasing cell size during development, PPARD activation, but not activation of PPARA or PPARG, increased cell size as well as the number of binucleated hPSC-CMs (**Figure 2E/F**). Together, our data show that PPAR signaling is not required for early cardiac specification. However, PPAR signaling in hPSC-CMs acts in an isoform-specific manner where PPARD activation results in enhanced sarcomere alignment, increased cell size and increased numbers of binucleated cells, all of which are changes consistent with enhanced cardiac maturation.

**Figure 2:**
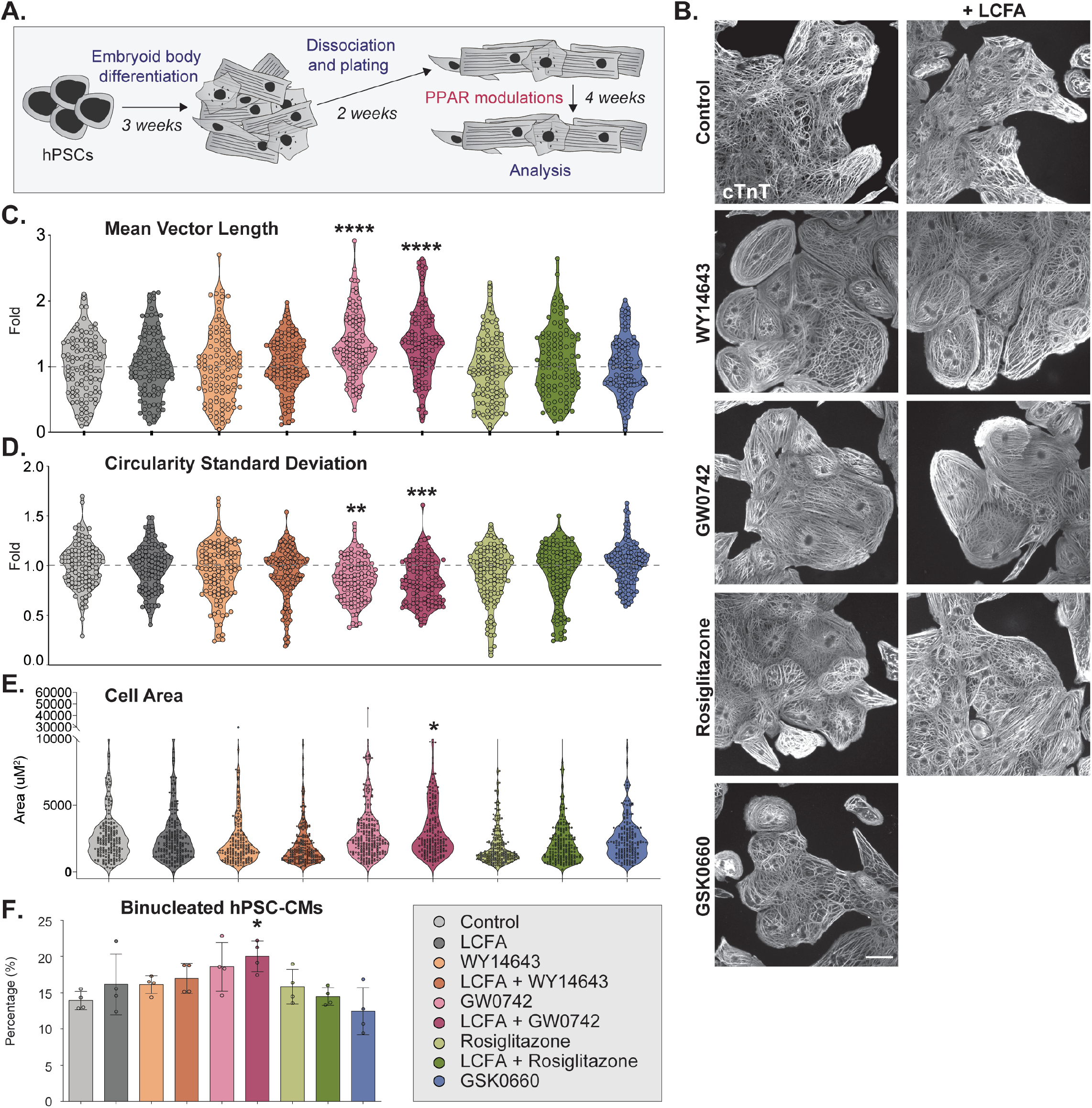
PPARD signaling activation induces changes in cell morphology, cell size and number of nuclei in hPSC-CMs. (**A**) Schematic of experimental outline. (**B**) IF analysis of hPSC-CMs 4 weeks after PPAR signaling modulations with antibodies against cardiac Troponin T (cTnT). Representative images are shown, scale bar = 50μm. (**C/D**) Mean Vector Length and Circularity Standard Deviation were quantified by MatFiber from IF analysis images for cTnT (n=3, >15 hPSC-CMs/biological replicate). (**E**) hPSC-CM cell area measured by cell surface area on images from IF analysis for alpha-actinin (n=3, >15 hPSC-CMs/biological replicate). (**F**) Percentage of binucleated hPSC-CMs were determined on images from IF analysis for alpha-actinin and DAPI stain (n=4, >250 hPSC-CMs/condition). Data represented as mean + SD. Statistics: One-way ANOVA and Tukey test for multiple comparisons (*p<.05; **p<.01); ***p<.001; **** p<.0001) relative to the control condition (far left).

### PPARD signaling induces a fatty acid oxidation transcriptional program

To further investigate the mechanisms underlying the changes observed in PPAR modulated hPSC-CMs, we performed RNA sequencing (RNAseq). In addition to the small molecules specific for all three PPAR isoforms, we added BSA-complexed long chain fatty acids (LCFAs: palmitic acid, oleic acid, linoleic acid) to address the effects of each PPAR isoform in the presence and absence of LCFA substrates. hPSC-CMs were differentiated and PPAR signaling induced/inhibited as described in Figure 2A. SIRPA+CD90-hPSC-CMs were FACS-purified 4 weeks after treatment for RNAseq and subsequent analysis on pure cardiac populations (Dubois et al., 2011; Elliott et al., 2011; Skelton et al., 2014). In agreement with the previous isoform-specific changes on cell morphology PPARD signaling modulations resulted in the greatest number of unique transcriptional changes in hPSC-CMs. Using a cutoff of p<0.05 we identified the highest number of differentially expressed genes relative to control in GW0742 treated (130 upregulated, 66 downregulated), LCFA + GW0742 treated (129 upregulated, 67 downregulated) and GSK0660 treated hPSC-CMs (107 upregulated, 109 downregulated), as well as a smaller number of differentially expressed genes in all of the other conditions (**Figure S3A**). KEGG pathway analysis revealed an upregulation of metabolism-related terms, such as ‘FA metabolism’, ‘Adipocytokine signaling pathway’ and ‘Peroxisomes’ in PPARD activated hPSC-CMs (**Figure 3A**). In accordance, inhibition of PPARD signaling results in a decrease of genes associated with the same KEGG pathways. Studies in both the mouse and PSC differentiations have found WNT signaling to be decreased during cardiac maturation (Buikema et al., 2020; Karakikes et al., 2014; Naito et al., 2006; Ueno et al., 2007; Uosaki et al., 2015). In line with these findings PPARD signaling induction results in a decrease of WNT pathway activators and an increase of WNT inhibitors (**Figure 3A/B**). Interestingly, we did not observe large changes in expression of genes encoding for sarcomere proteins, cardiac conduction calcium signaling components or cardiac transcription factors after PPARD activation (**Figure S3B**). However, PPARD activation upregulated many key genes involved in LCFA metabolism. These include transporters such as *CD36* (4.82-fold), which encodes the primary LCFA plasma membrane transporter and *CPT1A/CPT1B* (2.64-fold/1.45-fold), coding for Carnitine Palmitoyltransferase I, which is localized in the outer mitochondrial membrane and controls the rate-limiting step for the shuttle of LCFA-acyl-carnitines across the outer mitochondrial membrane (Figure 3B). They further include FAO enzymes such as *ACADVL* (2.31-fold), which codes for VLCAD, the enzyme that catalyzes the first step in LCFA beta-oxidation and *HADHA* (2.46-fold) and *HADHB* (2.73-fold), which catalyze the last three reactions in mitochondrial FAO (**Figure 3B**). Interestingly, *PDK4* was strongly upregulated (6.89-fold) by PPARd signaling activation. PDK4 inhibits glycolysis-dependent oxidative phosphorylation (OXPHOS) through the inactivation of Pyruvate Dehydrogenase, which has been demonstrated to be critical for cardiac maturation both *in vivo* and *in vitro* (Nakano et al., 2017). Additional candidates induced by PPARD signaling include *ACSL1*, which is involved in forming acyl-LCFAs, *MLYCD*, which catalyzes the conversion of malonyl-CoA to acetyl-CoA, and *ANGPTL4*, which is involved in lipoprotein lipase inhibition and triglyceride clearance as well as several other important FAO enzymes (**Figure 3B**). Previous work had suggested that the addition of LCFAs alone could enhance FAO in hPSC-CMs (Yang et al., 2019). In accordance with this, we observed enhanced expression of FAO genes 4 week after LCFAs supplementation (*CPT1A*: p=3.42E-01, *ACADVL*: p =5.30E-01, *PDK4*: p= 3.48E-01), however, this increase was lower than the combination of LCFAs with PPARD signaling activation. To assess whether PPARD signaling was essential for the induction of the FAO transcriptional program in hPSC-CMs, we analyzed gene expression after PPARD inhibition. Several of the key metabolism regulators including *CD36, CPT1A, ASCL1, CPT1B, HADHA, HADHB* and *PDK4* indeed showed significantly decreased expression compared to control hPSC-CMs, suggesting some endogenous activity of PPARD signaling in control conditions. We have made the complete gene expression data set available for exploration through the publicly accessible and searchable gene expression portal established as a resource for this project (https://maayanlab.cloud/dubois/).

**Figure 3:**
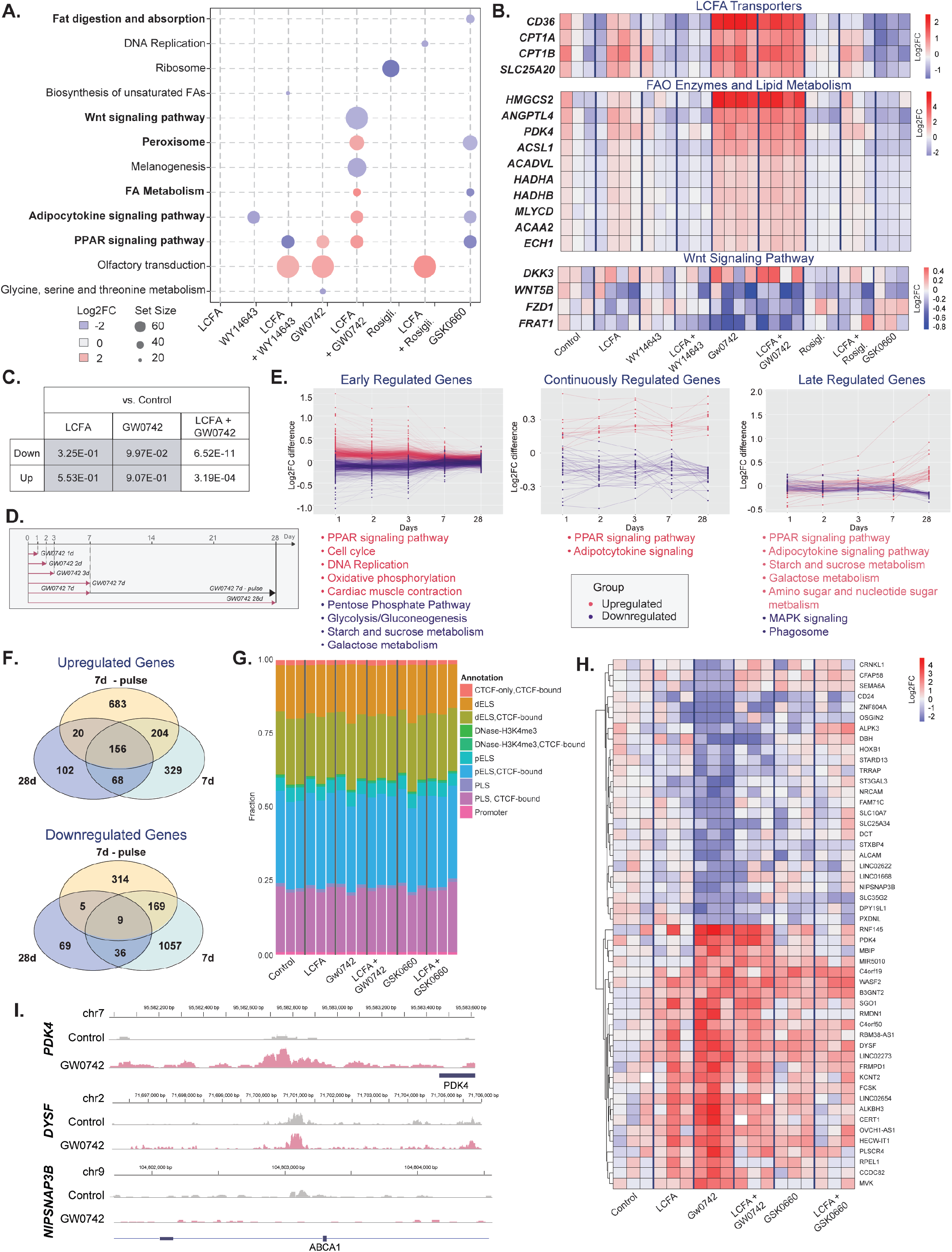
PPARD signaling activates the FAO transcriptional program in hPSC-CMs. (**A**) Comparison of KEGG pathways differentially regulated in PPAR modulated hPSC-CMs compared to control 4 weeks after continuous PPAR modulation (Set size – number of genes from KEGG pathway in dataset). (**B**) Expression of key candidates involved the FAO pathway: CD36, the primary LCFA plasma membrane transporter, CPT1A, a mitochondrial membrane LCFA transporter, ACADVL, the rate-limiting enzyme of mitochondrial FAO, and PDK4, an inhibitor of glycolysis-dependent OXPHOS, HADHA and HADHB, which catalyzes the last three reactions of FAO. DKK3, WNT5B, FZD1 and FRAT1 are regulatory components of the WNT pathway. (**C**) Comparison of genes differentially expressed in PPARD-modulated hPSC-CMs compared to control hPSC-CMs with gene sets of differentially expressed genes between D27 hPSC-CMs and the adult heart (Pavlovic et al., 2018). (**D**) Schematic of PPARD manipulations and hPSC-CM harvest timepoints. (**E**) Temporal analysis of gene expression changes upon PPARD induction (GW0742) for 1,2,3, and 7 days and 4 weeks revealed genes regulated significantly only within days 1, 2, 3 and 7 (left), genes that were continuously changed at all time points (middle), and genes regulated late during maturation (right). (**F**) Venn Diagram of upregulated (upper panel) and downregulated (lower panel) genes in hPSC-CMs after 7 days, a 7 day pulse and 28 days of PPARD activation. (**G**) Stacked bar chart of the quantities of promoter and enhance associated peaks identified by ATACseq across the 6 conditions assayed. (**H**) Heatmap of 50 genes with the greatest differential chromatin accessibility (25 most increased accessibility; 25 most decreased accessibility) between control and GW0742-treated hPSC-CMs displayed across the 6 conditions assayed. (**I**) ATACseq tracks for *PDK4, DYSF* and *NIPSNAP3B* displaying differential chromatin accessibility in promoter and enhancer regions in control and GW0742-treated hPSC-CMs. Statistics: DESeq2 was used to normalize read counts and determine differentially expressed genes with p<0.05 relative to the control condition (far left).

In summary, we show that the PPAR transcription factors act in an isoform-specific manner to activate gene regulatory networks involved in FAO, while simultaneously inhibiting glucose-dependent energy metabolism in hPSC-CMs. We further demonstrate that inhibition of PPARD results in inverse changes of gene expression, thus suggesting a direct role for PPARD signaling in regulating metabolic pathways in hPSC-CMs.

### PPARD activation results in sequential changes in gene expression in hPSC-CMs

Having identified the PPARD-mediated increase in transcription of genes required for oxidative metabolism in hPSC-CMs, we next determined whether these changes were indicative of metabolic maturation using comparative gene expression analysis. To support the hypothesis that PPARD activation drives hPSC-CMs along a developmentally relevant trajectory, we leveraged existing gene expression datasets that compared day 27 hPSC-CMs to the adult human heart (**Figure 3C**)(Pavlovic et al., 2018). We correlated the upregulated (indicative of enhanced maturation) and downregulated (indicative of decreased maturation) genes from the D27 hPSC-CM-with-adult heart comparison to the differentially expressed genes after PPARD perturbations. Interestingly, LCFA + GW0742 treatment was the only condition for which the differentially expressed genes exhibited a significant correlation with the genes that were upregulated (p= 6.52E-11) or downregulated (p= 3.19E-04) in the D27 hPSC-CM-with-adult heart. This unbiased analysis suggests that PPARD activation induces a gene expression profile that more closely correlates with mature CMs compared to any other culture condition.

Next, we investigated the temporal mechanisms by which PPARD activation induces FAO in hPSC-CMs. PPARs are transcription factors and we expected changes in gene expression to occur shortly after induction in hPSC-CMs, as well as potentially long-term changes reflecting a change in cell state. To distinguish these mechanisms we interrogated genes regulated shortly after PPARD activation/inhibition and genes differentially expressed after the 4 weeks of treatment as a consequence of changes over time in FACS-isolated hPSC-CMs (SIRPA+CD90-) and performed RNAseq (**Figure 3D**). In line with our hypothesis, expression analysis at the different times after PPARD activation identified groups of genes that were differentially regulated across time. These groups were i) Genes that were only changed during the first 7 days of treatment (early regulation genes, 695 upregulated and 1160 downregulated), ii) Genes that remained differentially expressed throughout the treatment timeline (continued regulation, 13 upregulated and 33 downregulated) and iii) Genes that were differentially regulated only at the end of the 4 week treatment (late regulation genes, 39 upregulated and 44 downregulated)(**Figure 3E**). Within all three subsets we identified candidates involved in various aspects of heart development (**Supplemental File 1**). To perform pathway analysis, we first collated the KEGG pathways identified from both up and downregulated genes in the GW0742, GW0742 1d, GW0742 2d, GW0742 3d and GW0742 7d treatment conditions (Table 1). The genes within the early, continued and late response groups were compared to the list of KEGG pathways and were extracted if there was a match. This match would indicate that the gene was a component of the listed KEGG pathway and allowed for the investigation of the pathways changing despite the small number of genes in the continued and late response groups. This analysis contributed to our understanding of underlying temporally-regulated mechanisms that are responsible for the PPARD-induced metabolic switch (**Figure 3E** and **Figure S4A**).

**Table 1.** Analysis of the differentially regulated KEGG pathways for genes within the early, continued and late response gene sets. The differentially regulated KEGG pathways for GW0742, GW0742 1d, GW0742 2d, GW0742 3d and GW0742 7d treatment conditions, relative to the 28-day control condition, were utilized to identify the genes classified as early, continued or later response genes belonging to these KEGG pathways.

As expected, the early transcriptional response to PPARD activation showed the greatest effect on differentially regulated pathways. For example, there was early upregulation of cell cycle and OXPHOS genes. Additionally, PPARA was downregulated, suggesting that PPARD activation may directly regulate the expression of the other PPAR isoforms. We further observed an early downregulation in genes related to glycolysis, galactose metabolism and pentose phosphate pathway. Interesting early candidates along these lines included *PFKM*, which catalyzes the rate-limiting step of glycolysis, and *PEX2*, which is a peroxisomal biogenesis factor (Shimozawa et al., 1992; Vora et al., 1980). Continuously up-regulated genes included *SLC25A20*, which is involved in the transport of acyl-carnitines into the mitochondrial matrix and *MLYCD*, a positive regulator of FAO. Within late response genes we identified candidates such as *HADHA* and *HADHB*, which catalyze the last three reactions in mitochondrial FAO (**Figure S4A**). Our data indicate that the majority of gene expression changes upon PPARD activation occur shortly after activation, with a subset of genes that are continuously changing over time and others that are only differentially regulated at later stages during the maturation process. An improved understanding of active pathways and detailed investigation of specific candidates over time after PPARD activation promises to further inform our understanding of the sequential mechanisms underlying the metabolic switch *in vitro*.

Given this dynamic temporal gene expression pattern we next addressed whether transient PPARD activation is sufficient to induce permanent gene expression changes in hPSC-CMs. To test this, we induced PPARD signaling for 7 days, then continued to culture the hPSC-CMs for an additional 21 days and performed RNAseq analysis on FACS-isolated cells (7d-pulse). The comparison of the differentially expressed genes in the 7d-pulse to those after 7 or 28 days of continued PPARD signaling confirmed the existence of a temporal dynamic after PPARD induction (**Figure 3F** and all differentially expressed genes summarized in **Supplemental File 2**). We identified many genes shared between the 7d-pulse and the 7d and 28d continuous induction (156 upregulated genes and 9 downregulated genes), indicating that a large number of PPARD-induced genes remain differentially expressed even in the absence of continued pathway activation. However, other differentially regulated genes did segregate according to the duration of PPARD induction, suggesting that PPARD signaling duration impacts the extent of transformation, with the long-term induction (4 weeks) leading to the most distinct gene expression changes. All of the gene expression data described here is included in the searchable gene expression database for further exploration of individual candidates of interest (https://maayanlab.cloud/dubois/).

The permanent changes in gene expression after transient PPARD induction suggested a potential regulation on the epigenome level. To address this we performed ATAC sequencing (ATACseq) after PPARD signaling activation and inhibition. When assaying across all of the 6 conditions measured (control +/- LCFA, GW0472 +/- LCFA, GSK0660 +/- LCFA), we did not observe significant differences in the overall amount of promoter and enhancer-associated peaks, as expected (**Figure 3G**). In concordance with the RNAseq data however, we found a segregation between control and PPARD-activated hPSC-CMs through Principal Component Analysis (PCA)(**Figure S4B**), suggesting that PPARD activation is able to modify chromatin accessibility in hPSC-CMs. We performed differential peak analysis to identify promoter or enhancer associated peaks with the biggest increases and decreases in chromatin accessibility between control and PPARD-activated hPSC-CMs and discovered that PPARD activation enriched a specific population of genes, many of which were shared in the LCFA + GW0742 condition (**Figure 3H**). For example, we identified an increase in the chromatin accessibility upstream of *PDK4*, correlating with increased *PDK4* gene expression uncovered earlier (**Figure 3I**). Similarly, *DYSF* and *KCNT2* both showed increased chromatin accessibility, where the former encodes a Ca^2+^ sensor involved in maintaining muscle integrity and the latter encodes an outward rectifying K^+^ channel (**Figure 3I** and **Figure S4C**)(Bhattacharjee et al., 2003; Han et al., 2007). We also identified genes with reduced chromatin accessibility in their regulatory domains, including for example *DYSF, NIPSNAP3B, PXDNL, ALCAM, SEMA6A, HOXB1* and *TRRAP* for which roles in cardiac development and maturation are just beginning to emerge and making them exciting to study in this context (**Figure 3H/I** and **Figure S4D**)(Péterfi et al., 2014; Xu et al., 2021).

In conclusion, our data illustrate that PPARD signaling activation in hPSC-CMs results in a transcriptional and epigenetic profile that more closely resembles that of mature heart tissue compared to control hPSC-CMs. Our data further demonstrate that PPARD activation causes sequential activation of the gene regulatory networks involved in establishing the FAO machinery in hPSC-CMs and that transient PPARD activation is able to induce permanent changes in the gene regulatory network driving FAO, albeit not to the same extent as continuous activation.

### PPARD activation increases *in vitro* LCFA uptake and processing

To determine if the increased expression of the FAO machinery after PPARD activation translates to augmented FAO in hPSC-CMs we investigated the effects of PPARD signaling on LCFA uptake and processing. We focused on transport proteins required for LCFA shuttling across the plasma and mitochondrial membranes and the enzymes catalyzing the beta-oxidation of LCFA in the mitochondria. CD36 is the primary LCFA transporter on the plasma membrane and is responsible for 70% of LCFA entry into the cell (Coburn et al., 2000; Glatz et al., 2016). In the absence of CD36, LCFAs are unable to enter the cell (Glatz et al., 2016). Furthermore, CD36 has recently been described as a marker for a subset of hPSC-CMs harboring more mature characteristics (Funakoshi et al., 2021; Poon et al., 2020). We performed flow cytometry analysis of hPSC-CMs treated with GW0742 (PPARD agonist) or GSK0660 (PPARD antagonist) for 4 weeks to quantify cell surface expression of CD36. Interestingly, control hPSC-CMs expressed low levels of CD36, suggesting that LCFA uptake may be limited with current differentiation strategies (**Figure 4A/B**). Consistent with the increased *CD36* gene expression, PPARD induction significantly increased CD36 cell surface expression by 13.36-fold (**Figure 4A/B**). In addition to serving as a substrate for FAO, LCFAs can act as PPAR ligands and LCFA supplementation has been previously shown to enhance hPSC-CM maturation (Yang et al., 2019). Consistent with these observations we also observed a 2.33-fold increased CD36 cell surface expression in LCFA-supplemented cultures. This increase was consistently lower than what can be achieved via PPARD activation, and is not dependent on LCFA concentration (**Figure 4B** and **Figure S5A**). To ensure that the effects of PPARD induction were reproducible across hPSC lines, we confirmed the ability of PPARD activation to increase CD36 cell surface expression in 2 additional hPSC lines (hiPSCs and hESCs)(**Figure S5B/C**). We next generated atrial hPSC-CMs to investigate the effect of PPARD activation on the FAO machinery in non-ventricular CMs. Using a differentiation protocol with retinoic acid (RA) supplementation during the mesoderm induction stage, we demonstrate that CD36 is also absent in atrial hPSC-CMs under baseline conditions, and that PPARD activation induces CD36 cell surface expression to similar extents as in ventricular hPSC-CMs, suggesting that this is a mechanism common to different cardiac cell types (**Figure 4C**)(Devalla et al., 2015; Lee et al., 2017; Zhang et al., 2011).

**Figure 4:**
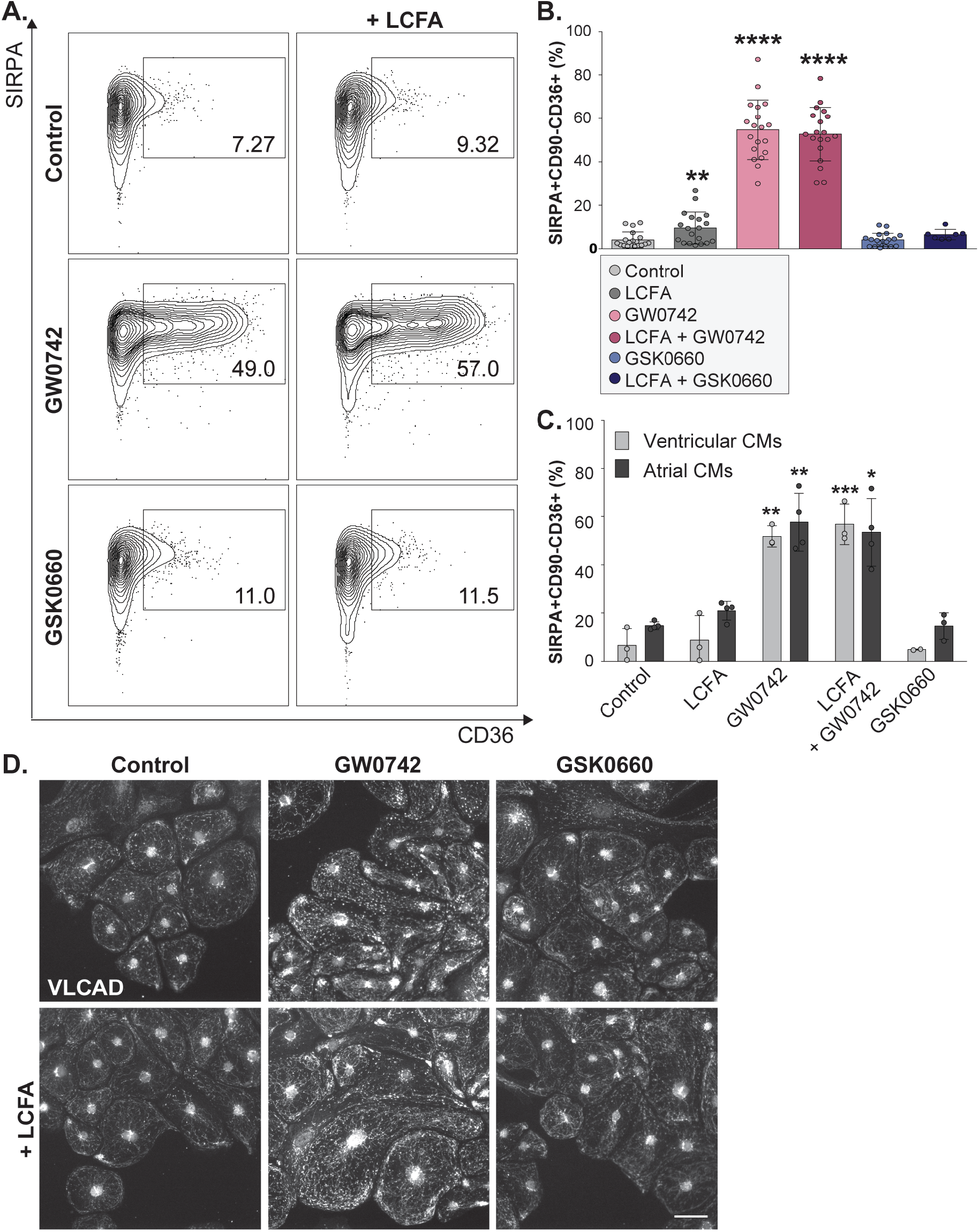
PPARD signaling enhances LCFA uptake and processing in hPSC-CMs. (**A**) Flow cytometry analysis of hPSC-CMs 4 weeks after continuous PPAR modulations. SIRPA marks hPSC-derived cardiomyocytes; CD36 is a key LCFA transporter expressed on the plasma membrane. (**B**) Quantification from (A), (n=7-19 biological replicates). **(C**) Flow cytometry analysis of CD36 expression after PPAR modulation in ventricular and atrial cardiomyocytes (n=2-4). (**D**) IF analysis for VLCAD, the rate-limiting enzyme of FAO, on hPSC-CMs 4 weeks after continuous PPAR modulation. Scale bar = 50μm. Data represented as mean + SD. Statistics: Student’s t-tests relative to the control condition (far left) (*p<.05; **p<.01; ***p<.001; **** p<.0001).

After crossing the cell membrane, LCFAs are activated to an acyl-CoA ester, enter the mitochondria via the carnitine shuttle and are subsequently processed by a series of FAO enzymes. In agreement with its transcriptional upregulation, PPARD activation increased VLCAD protein expression in hPSC-CMs and led to a more widespread expression pattern across the cells (**Figure 4D**). Moreover, FAO, particularly for LCFAs with greater than 20 carbons occurs in both mitochondria and peroxisomes. To assess the effects of PPARD activation on peroxisomes we stained hPSC-CMs with antibodies against PMP70, a peroxisomal protein. We detected significantly increased peroxisomal content after PPARD activation, as shown by IF and quantified by flow cytometry (**Figure S5D/E**).

In conclusion, we show that PPARD activation induces the FAO machinery in hPSC-CMs by efficiently increasing LCFA transporters and key FAO enzymes, as well as the total peroxisome content, suggesting that this represents a mechanism to robustly enhance both the uptake and utilization of LCFAs in a variety of hPSC-derived cardiac cell types.

### PPARD activation enhances *in vitro* OXPHOS

CM-restricted PPARD knockout mouse models have shown that PPARD signaling regulates mitochondrial biogenesis through Ppargc1a, the PPAR signaling pathway transcriptional coactivator which we find upregulated after PPARD activation (**Figure 3B** and **Figure S6C**)(Liu et al., 2011). Increased mitochondrial content is a hallmark of CM maturation, as mature CMs require more mitochondria in close proximity to the contractile apparatus to rapidly provide the energy required for contraction. To investigate changes in mitochondrial content and distribution after PPARD manipulation, we stained hPSC-CMs with MitoTracker Deep Red dye 4 weeks after treatment. IF analysis illustrates that PPARD induction leads to an increase in mitochondrial content and a greater distribution of mitochondria from the perinuclear localization typically observed in control hPSC-CMs to a more widespread distribution across the entire cell and specifically in between sarcomere structures (**Figure 5A**). Quantification of the intracellular MitoTracker Deep Red fluorescence was performed via flow cytometry, demonstrating a 1.16- and 1.25-fold increase in mitochondrial content in hPSC-CMs after PPARD activation -/+ LCFAs (**Figure 5B/C**). To assess the mitochondrial structure, which becomes increasingly complex during cardiac maturation, we performed transmission electron microscopy (TEM) analysis after PPARD activation or inhibition. Morphometric quantification of TEM images revealed that both PPARD activation and LCFA supplementation increased mitochondrial surface area, with the greatest change observed in PPARD activation in the presence of LCFAs (**Figure 5D/E**). Similarly, only LCFA in combination with GW0742 significantly increased both mitochondrial length and width (Figure 5F and Figure S6A). Lastly, mitochondrial cristae organization, a hallmark of mature and highly active mitochondria, was also improved in PPARD-induced hPSC-CMs (**Figure S6B**). In line with these observations, we found *MFN2*, a key driver of mitochondrial fusion, to be upregulated in PPARD-activated hPSC-CMs, while *DNM1*, a regulator of mitochondrial fission was unchanged (**Figure S6C**).

**Figure 5:**
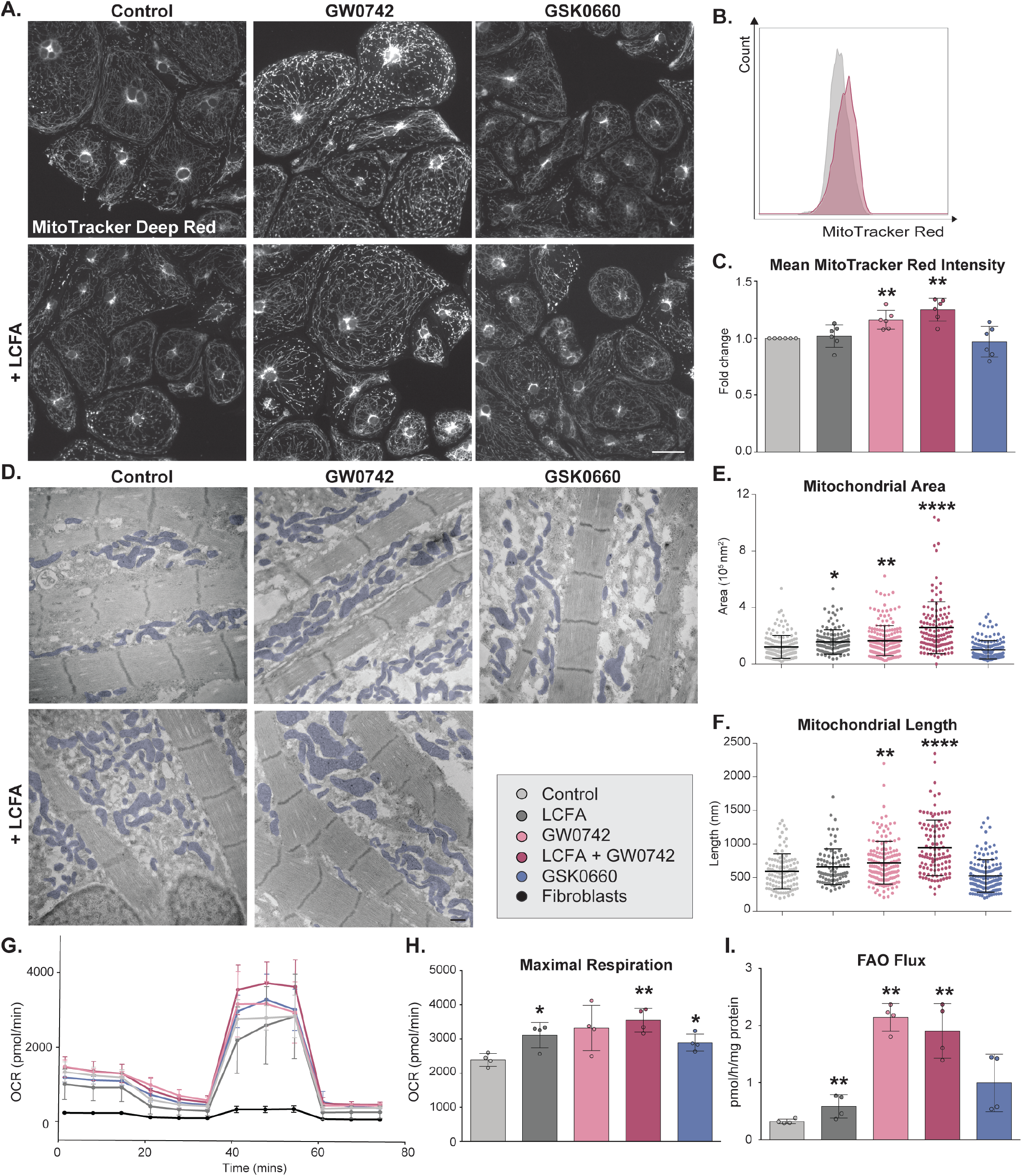
PPARD signaling induces metabolic maturation in hPSC-CMs. (**A**) MitoTracker Deep Red dye analysis in hPSC-CMs 4 weeks after continuous PPARD modulations. Scale bar = 50μm. (**B/C**) Quantification of mitochondrial content by flow cytometry analysis for MitoTracker Deep Red. Representative flow cytometry plots from Control (grey) and LCFA + GW0742 (purple) treated hPSC-CMs (B) and quantification across all conditions (**C**) (n=6). (**D**) Transmission electron microscopy analysis of hPSC-CMs 4 weeks after continuous PPARD modulations. Mitochondria are pseudo-colored in blue. Scale bar = 500nm. (**E/F**) Quantification of mitochondrial area and length on TEM images using imageJ. Data collected from 5-10 images per conditions, with a minimum of 10 mitochondria per image. (**G/H**) Oxygen consumption rate (OCR) in hPSC-CMs and hPSC-derived fibroblast cells (black) 4 weeks after continuous PPARD modulations was measured using a Seahorse analyzer to calculate maximal respiration (n=4; n=5-8 technical replicates/ condition). (**I**) FAO Flux assay in hPSC-CMs 4 weeks after PPARD modulation (n=4). Data represented as mean + SD. Statistics: (**C, H, I**) Student’s t-tests relative to the control condition (far left) (*p<.05; **p<.01; ***p<.001; **** p<.0001). (**E, F**) One-way ANOVA and Tukey test for multiple comparisons (*p<.05; **p<.01); ***p<.001; **** p<.0001) relative to the control condition (far left).

To determine if the increase in mitochondria number and mitochondrial size in PPARD-activated hPSC-CMs resulted in greater metabolic capacity we quantified mitochondrial oxidative phosphorylation (OXPHOS). Basal respiratory capacity was similar between hPSC-CMs treated with GW0742 or GSK0660 for 4 weeks (**Figure S6D**). However, PPARD-activated hPSC-CMs exhibited a 1.82-fold increase in maximal respiration (rate of oxygen consumption in the presence of the mitochondrial uncoupler FCCP, which eliminates the proton gradient across the inner mitochondrial matrix, minus the non-mitochondrial oxygen consumption), and a 1.42-fold increase in the mitochondrial spare capacity (difference in oxygen consumption between maximal and basal respiration) indicative of an augmented capacity to generate ATP upon increased metabolic demand (**Figure 5G/H** and **Figure S6D**). Interestingly, increase in the maximal respiration and the spare capacity was also observed to lower rates in hPSC-CMs treated with LCFAs alone or GSK0660 (maximal respiration: LCFAs: 1.30-fold, GSK0660: 1,21-fold; Spare Capacity: LCFAs: 1.43-fold, GSK0660: 1.40-fold). While informative on overall ATP generation, this analysis does not distinguish FAO-related from non FAO-related oxidative reactions,

For the highly metabolically active adult heart, metabolic flexibility with respect to substrate utilization is critical to ensure the organ’s continuous function. LCFAs are an advantageous substrate as beta-oxidation produces more ATP per molecule than glucose metabolism, and active FAO is characteristic of a healthy heart (Lopaschuk and Jaswal, 2010). OXPHOS measured in the Seahorse Mito Stress Test is an incomplete metric of active FAO, as it measures both glucose- and LCFA-derived oxidation. We therefore addressed the extent of FAO more specifically by quantifying the rate of radiolabeled [9,10-3H]-Palmitate oxidation, which constitutes a more definitive assay for FAO (Doulias et al., 2013). In correlation with our previous results, PPARD activation, led to a significant increase in FAO flux compared to control hPSC-CMs (6.70-fold and 6.00-fold increase for GW0742 and GW0742 + LCFA respectively), while LCFA supplementation alone contributed to a 2.43-fold increase (**Figure 5I**).

Collectively our data demonstrate that PPARD activation promotes FAO by improving the ability of hPSC-CMs to uptake and metabolize LCFAs *in vitro*, while simultaneously limiting the contribution of glucose-dependent ATP production. PPARD signaling activation thus induces the metabolic switch *in vitro* that is characteristic of the metabolic maturation observed during *in vivo* heart development.

### PPARD signaling activation enhances electrophysiological and contractile maturation in hPSC-CMs

Continuous myocardial compaction and the systemic pressure of blood flow result in a continuous increase in contractility of the heart over the course of development, which coincides with the increase in PPAR signaling activity starting at midgestation (Uosaki et al., 2015). In line with a potential role for PPAR signaling for cardiac contractility, cardiac-specific PPARD deficiency in mice leads to cardiomyopathies and contractile defects (Cheng et al., 2004). Additionally, enhanced metabolic activity via LCFA addition in hPSC-CMs was shown to increase contractility *in vitro* (Feyen et al., 2020). To assess cardiac electrophysiology and contractility after PPARD activation/ inhibition we performed action potential (AP) measurements, Ca^2+^ transient (CaT) analysis and contractility measurements in engineered heart tissues (EHTs). Using the experimental protocol depicted in Figure 2A, hPSC-CMs were dissociated and plated as single cells 4 weeks after PPARD activation/ inhibition and paced at 0.5Hz for all of the measurements. CaT analysis demonstrated no differences in amplitude, but significant differences in the 20th and 50th percentiles of the intracellular calcium transient duration (GW0742: 1.1-fold CaD50; 1.24-fold CaD20; GW0742 + LCFA: 1.10-fold CaD50, 1.17-fold CaD20) and decreased by 0.91-fold in PPARD-inhibited cells (**Figure 6A**). Increased CaT duration is indicative of increased availability of Ca^2+^ for contraction (see correlative EHT data later in this section). PPARD activation further resulted in increased action potential (AP) durations (GW0742: 1.24-fold APD50; 1.25-fold APD20; GW0742 + LCFA: 1.25-fold APD50, 1.23-fold APD20) and PPARD inhibition in turn resulted in an APD50 reduction to 0.89-fold relative to control hPSC-CMs, with no change in amplitude in any of the conditions (**Figure 6B**). This increase in APD may be in part due to increased duration of Ca^2+^ flux during the plateau phase of the cardiac AP, establishing strong correlation between the different measurements in PPARD manipulated hPSC-CMs. In line with these findings, previous maturation protocols have resulted in similar changes to the AP and in CaT (Correia et al., 2017, 2018; Tiburcy et al., 2017). To study the effect of PPAR modulation on cardiac function in a group of connected cells we performed multi-electrode array (MEA) analysis on FACS-isolated hPSC-CMs, and found no changes in field potential duration and conduction velocity after PPARD activation or inhibition (**Figure 6C**). This is perhaps unsurprising given the non-paced conditions used for this data acquisition. Collectively, our electrophysiology analysis suggests that PPARD activation improves electrophysiological maturation in hPSC-CMs.

**Figure 6:**
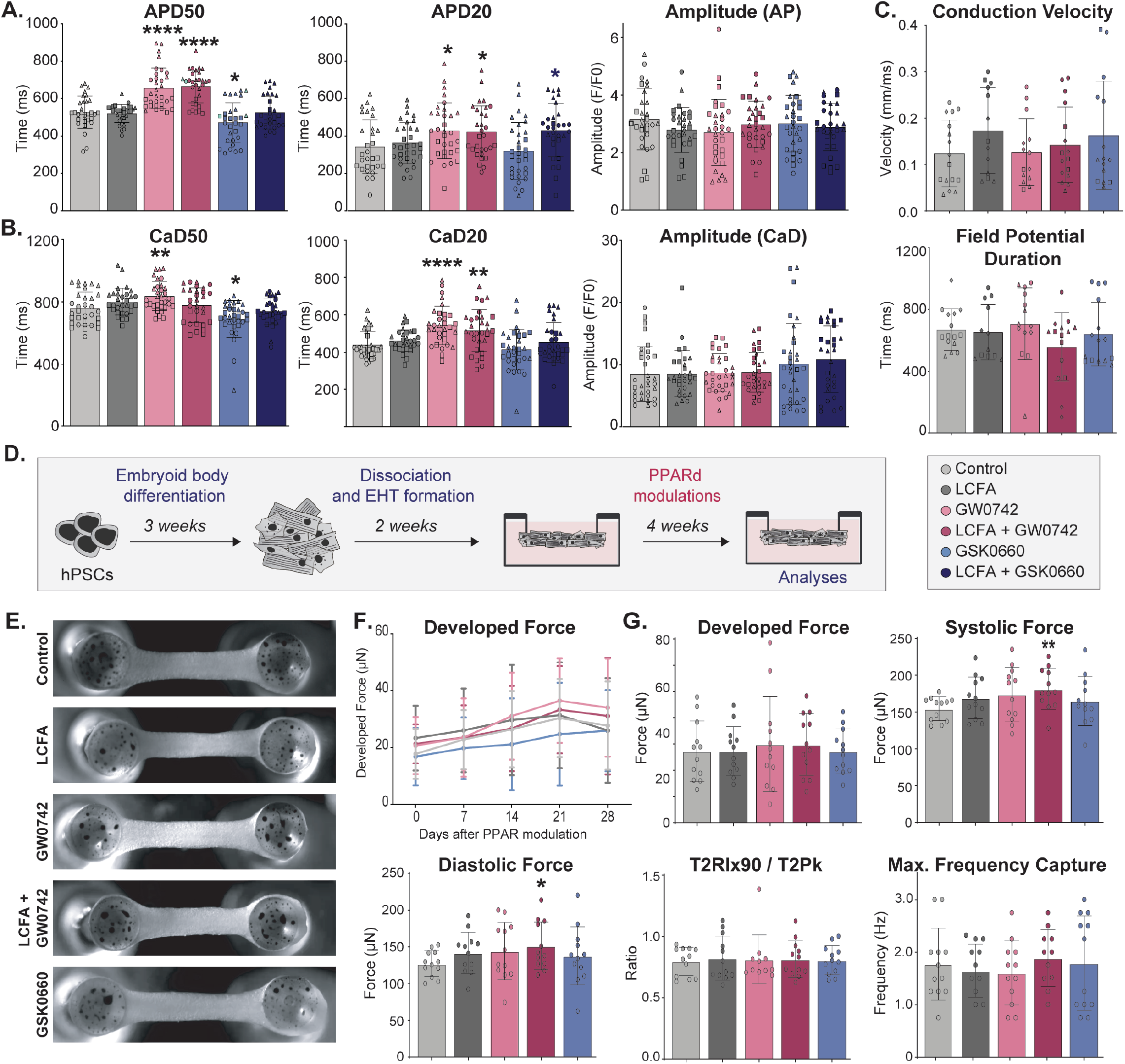
PPARD signaling improves hPSC-CM electrophysiological and contractile maturation. (**A**) Action potential (AP) measurements on single hPSC-CMs 4 weeks after continuous PPARD activation/inhibition. hPSC-CMs were paced at 0.5 Hz (n=3 biological replicates, 4-10 hPSC-CMs measured/ condition). (**B**) Calcium transient analysis on single hPSC-CMs 4 weeks after continuous PPARD activation/inhibition. hPSC-CMs were paced at 0.5 Hz (n=3 biological replicates; 4-10 hPSC-CMs measured/condition). (**C**) Multi-electrode array (MEA) analysis of FACS-isolated hPSC-CMs 4 weeks after continuous PPARD activation/inhibition (n=4 biological replicates). (**D**) Schematic of experimental outline for Engineered Heart Tissue (EHT) formation. (**E**) Brightfield images of EHTs at 4 weeks after PPAR modulation. (**F**) Contractile force measurements in spontaneously-contracting EHTs during the 4 weeks of continuous PPARD modulations. (**G**) Analyses of EHTs 4 weeks after continuous PPARD modulations. EHTs were paced at 1.0 Hz (n=4; 3 EHTs/ biological replicate). Biological replicates are identified by same shape of respective data points. Data represented as mean + SD. Statistics: Student’s t-tests relative to the control condition (far left)(*p<.05; **p<.01; ***p<.001; **** p<.0001).

Finally, we explored the effect of PPARD modulation in 3D engineered heart tissues (EHTs), which represent the most physiologically relevant model system for measuring *in vitro* cardiac contractility (Mannhardt et al., 2017; Ruan et al., 2016; Schaaf et al., 2014). Culturing hPSC-CMs in EHTs has been shown to induce hPSC-CM structural maturation as characterized by aligned hPSC-CMs, tissue compaction and increasing developed force over time (Lemoine et al., 2017; Ronaldson-Bouchard et al., 2018; Tiburcy et al., 2011; Turnbull et al., 2014). EHTs were generated at day 20 of differentiation, allowed to compact for two weeks and treated with small molecule agonist and antagonists for 4 weeks (**Figure 6D**). EHTs in all conditions showed the anticipated increase in developed force over the 4 weeks, with an average 1.48-fold increase across all EHTs, and no difference in tissue compaction or gross EHT morphology (**Figure 6E/F** and **Supplemental File 3**). Neither PPARD activation nor inhibition affected the spontaneous developed force in EHTs between weeks 1-4 of treatment (**Figure 6F**). At a paced beat frequency of 1.0 Hz, the developed force, T2Rlx90 / T2Pk ratio and maximum capture frequency across EHTs remained unchanged across treatment conditions (**Figure 6G**). However, both systolic and diastolic forces were significantly increased by 1.17 and 1.19-fold respectively 4 weeks after LCFA + GW0742 treatment (**Figure 6G)**. These data support the changes observed in the AP and CaT measurements and demonstrate that PPARD signaling activation results in enhanced contractility in hPSC-CMs.

### Transient lactate exposure induces permanent gene expression changes in hPSC-CMs

In the context of exploring metabolic substrates and cardiac maturation more broadly we next turned to investigating the effect of lactate substrate on hPSC-CMs. Prior to switching to FAO postnatally, the developing heart relies on glucose and lactate as its main metabolic substrates (Burd et al., 1975; Lopaschuk and Jaswal, 2010; Neely and Morgan, 1974; Piquereau and Ventura-Clapier, 2018; Werner and Sicard, 1987). *In vitro*, the ability of hPSC-CMs to metabolize lactate as a substrate has been leveraged into widely-adopted hPSC-CM enrichment protocols (Tohyama et al., 2013). This is based on the ability of hPSC-CMs, but not other cell types generated during hPSC differentiation, to survive in glucose-free medium supplemented with only lactate for a limited amount of time. The effects of inducing lactate metabolism however have not been investigated in depth to date. In an unbiased approach we compared the transcriptome of control and lactate-treated hPSC-CMs 4 weeks after lactate selection. Surprisingly, transient lactate treatment was sufficient to result in extensive long-term transcriptional changes in hPSC-CMs 4 weeks after the last exposure to lactate. KEGG pathway analyses revealed downregulation of genes involved in processes such as ‘Oxidative Phosphorylation’ and ‘Galactose Metabolism’ (**Figure 7A/B**). Intriguingly, the pentose phosphate pathway was also downregulated upon lactate exposure, a pathway that had previously been described to enhance hPSC-CM maturation when inhibited (Nakano et al., 2017). Amongst the changes in metabolism-related genes, we identified several candidates that are involved in reducing lactate sensitivity and processing ability (**Figure 7C**). For example, *SLC16A3* encodes MCT4, which transports lactate across the plasma membrane, **SLC2A1** which encodes the glucose transporter GLUT1 and *LDHA, PDHA1, HK1* and *HK2*, which encode additional components involved in lactate turnover and glucose metabolism. Collectively these genes showed decreased expression, indicating that a transient pulse of lactate is sufficient to induce a long-term decreased sensitivity to the substrate, potentially recapitulating a decrease in lactate metabolism during cardiac development *in vivo*. Similar to PPARD induction, transient lactate exposure further leads to increased expression of WNT inhibitors such as *SFRP1* and *SRFP2*, and a decrease in *WNT11*, as well as decrease in proliferation markers (*CCND2, IGF2*)(**Figure 7C**). This is in line with other studies suggesting that WNT inhibition and decrease in proliferation are critical for cardiac maturation. Lastly, we observed increased expression of genes underlying cardiac contractility (*TNNT2, MYLK, MYOM3*) and calcium signaling (*CACNA1C, CASQ2*)(**Figure 7C**). These analyses suggest that the transient lactate exposure commonly used in the lactate selection protocol results in lasting gene expression changes in hPSC-CMs, potentially driving contractile maturation however without enhancing metabolic maturation.

**Figure 7:**
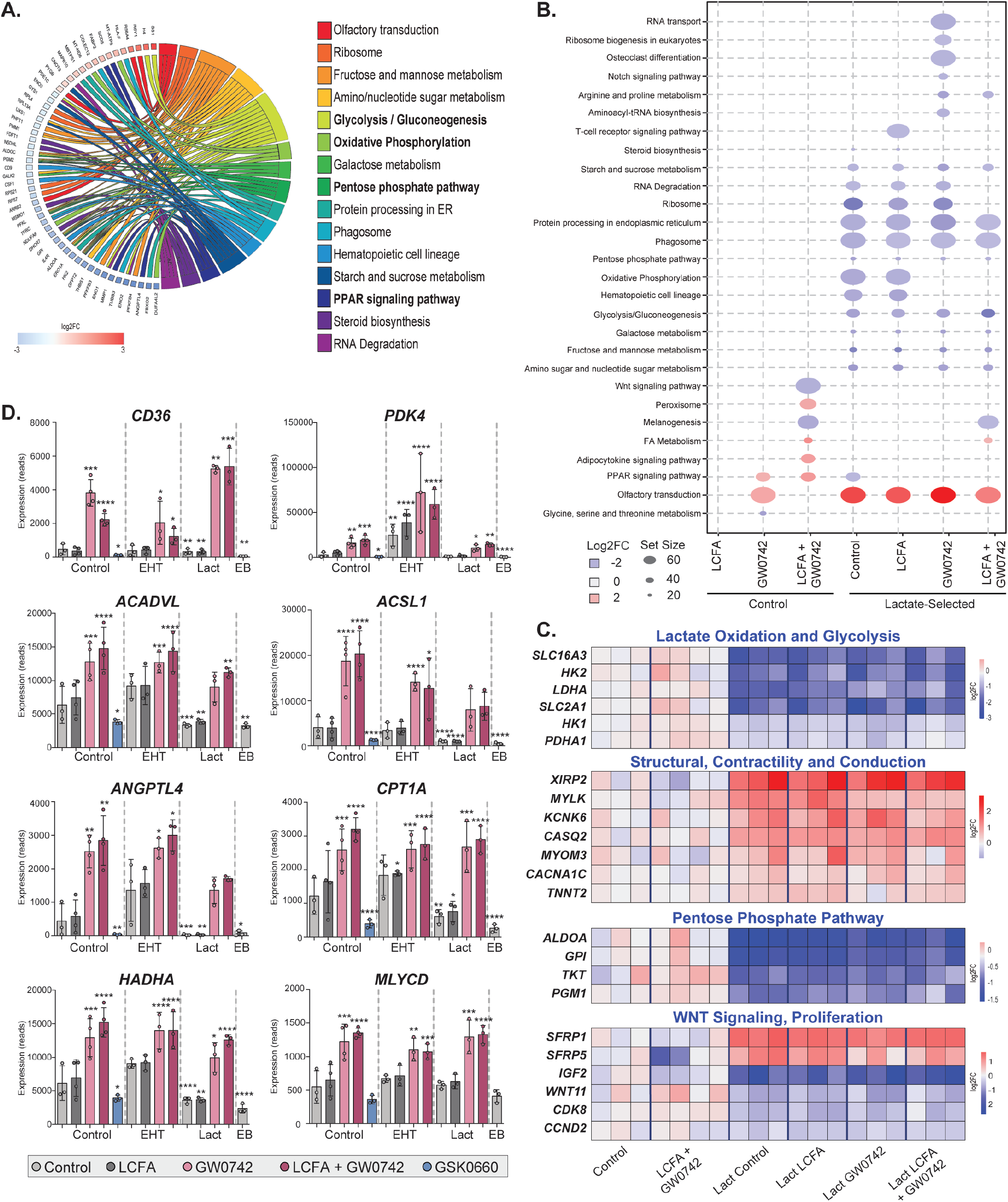
Transient lactate exposure shows long-term effects on cardiac maturation and PPARD activation induces the metabolic switch across multiple culture systems. (**A**) Chord plot illustration of differentially regulated KEGG pathways in hPSC-CMs transiently exposed to lactate compared to control hPSC-CMs. (**B**) KEGG pathways analysis and comparisons between hPSC-CMs after lactate selection and 4 weeks of continuous PPARD modulations relative to the control condition (far left). (**C**) Gene expression analysis (RNAseq data, n=3) in hPSC-CMs transiently exposed to lactate followed by PPARD modulation (4 weeks). (**D**) Gene expression analysis (RNAseq, n=3) of PPARD modulated FACS-isolated hPSC-CMs from regular 2D monolayers (Control), EHTs, lactate selection or of age-matched embryoid bodies (EB). Data represented as mean + SD. Statistics: DESeq2 was used to normalize read counts and determine differentially expressed genes with p<0.05 relative to the control condition (far left).

### PPARD induction induces a metabolic switch across multiple cell culture formats

Given the many different protocols used to generate, purify and analyze hPSC-CMs, and our data suggesting a lasting effect after lactate purification we next tested whether PPARD can effectively induce the metabolic switch in lactate-selected and EHT hPSC-CMs. To assess this, we induced PPARD signaling in lactate-purified hPSC-CMs and in EHTs for 4 weeks and performed RNAseq of FACS-isolated hPSC-CMs (SIRPA+CD90-). When combined with PPARD signaling activation, both lactate-selected hPSC-CMs and hPSC-CMs from EHTs showed a similar metabolism-focused transcriptional response as previously observed in 2D monolayer cells (**Figure 7C/D** and **Figure S7**). Lactate treatment or culture as EHTs followed by PPARD activation increased the expression of LCFA transporters (*CD36* and *CPT1A*), key enzymes supporting FAO (*ACADVL, ANGPTL4, HADHA, HADHB, ACSL1* and *PDK4*) and the ‘FA metabolism’, ‘Oxidative Phosphorylation’ and ‘Peroxisome’ KEGG pathways. Interestingly, both baseline expression (control, untreated) and the extent of differential gene expression after PPARD induction varied between culture formats, suggesting that there may be different metabolic activity in different formats, and that cells in such different formats may be differentially able to induce a metabolic switch after PPARD induction. This is supported by studies in EHTs demonstrating that larger tissue formats induce both contractile and metabolic maturation in the absence of any additional modifications (Ronaldson-Bouchard et al., 2018; Ulmer and Eschenhagen, 2020). In summary, transcriptional profiling across hPSC-CM culture formats suggests that PPARD activation is a reliable approach to induce the metabolic switch and to enhance FAO across multiple different culture formats, providing new opportunities for incorporating metabolic maturation broadly across diverse cardiovascular studies with hPSC-CMs and generating new multi-parameter strategies for metabolic and contractile maturation *in vitro*.

## DISCUSSION

The goal of this study was to improve the *in vitro* maturation of hPSC-CMs by leveraging some of the key mechanisms guiding heart development *in vivo* that are not recapitulated in current differentiation protocols. Due to the pressing need to generate more mature PSC-derived cell types in culture, other groups have pursued similar approaches, leading to remarkable progress in the field to date. These include, but are not limited to prolonged time in culture (DeLaughter et al., 2016; Kamakura et al., 2013), manipulation of signaling pathways and metabolism (Correia et al., 2017; Feyen et al., 2020; Funakoshi et al., 2021; Hu et al., 2018; Kosmidis et al., 2015; Murphy et al., 2021; Nakano et al., 2017; Parikh et al., 2017; Poon et al., 2015; Yang et al., 2019, 2014b), combinations of cardiovascular cell types (Pasquier et al., 2017; Vollert et al., 2013), tissue engineering, enhanced load, electrical stimulation (Correia et al., 2018; Giacomelli et al., 2020; Leonard et al., 2018; Li et al., 2018; Nunes et al., 2013; Ronaldson-Bouchard et al., 2018; Shadrin et al., 2017; Thavandiran et al., 2013; Tiburcy et al., 2017) and transplantation into *in vivo* model systems (Chong et al., 2014; Gao et al., 2018; Ogasawara et al., 2017; Shadrin et al., 2017). Despite these important advances, *in vitro* maturation of hPSC-CMs remains a challenge, and the field still lacks easy and broadly applicable strategies culminating in long-term enhanced maturation beyond early fetal stages.

Here we focused on the signaling pathways that are activated at the early stages of heart development, with a specific interest in those that are thought to regulate metabolism. We hypothesized that an incomplete recapitulation of these developmental mechanisms constitutes a limiting factor behind the immature hPSC-CMs generated through existing differentiation protocols. Accordingly, we focused on the PPAR signaling pathway due to its overall importance for the heart, and discovered that modulating PPAR signaling enhances maturation of hPSC-CMs by inducing the metabolic switch that occurs during development of the heart and shortly after birth.

PPARs, which comprise the isoforms PPARA, PPARD and PPARG, are nuclear receptors that act as transcription factors to regulate the expression of genes involved in processes such as FAO, proliferation and the immune response amongst others, with receptor levels varying across organs (Lee and Kim, 2015; Lopaschuk and Jaswal, 2010). Previous work in hPSC-CMs has not yet elucidated any distinct roles for the individual isoforms in hPSC-CMs and has rather focused on the most prominent isoform, PPARA (Funakoshi et al., 2021; Gentillon et al., 2019; Murphy et al., 2021; Poon et al., 2015; Uosaki et al., 2015). However, despite a large overlap of target genes between PPAR isoforms, their gene expression pattern and the distinctly different mouse KO phenotypes suggest that isoform-specific mechanisms are at play in different cells and tissues, and at different times during development and homeostasis. By activating and inhibiting each PPAR isoform independently, we discovered that PPARD signaling activation has specific transcriptional, structural and functional consequences in hPSC-CMs. PPARD signaling activation improves myofibril organization, cell size and binucleation while also inducing metabolic, electrophysiological and contractile maturation. We further uncovered an important role for PPARA during both cardiac differentiation and maturation, suggesting that PPARA is both active and required for hPSC-CM differentiation. However, additional activation using PPARA agonists did not resulted in significant changes in hPSC-CMs in our system. Other studies have utilized PPARA-specific agonists, either in combination with other modifications (LCFA supplementation and T3), or at a range of concentrations up to 20-fold higher than the ones used here and have indeed seen maturation effects, indicating that future strategies might combine these different modulations for potentially even greater maturation effects (Funakoshi et al., 2021; Gentillon et al., 2019; Poon et al., 2015). Lastly, PPARG did not impact *in vitro* maturation, in line with it being predominantly found in adipose tissue and being the lowest expressed isoform in the heart (Braissant and Wahli, 1998; Kliewer et al., 1994; Lee and Kim, 2015). Such isoform-specific roles are not surprising, given what is known about the function of PPARs *in vivo*, where genetic loss-of-function studies support an important role for both PPARA and PPARD in the heart (Cheng et al., 2004; Liu et al., 2011). Both receptors are involved in the regulation of FAO in mitochondria and peroxisomes, as well as in regulating the fatty acid import machinery. However, PPARA KO mice are viable throughout development, with lower FAO rates and a higher sensitivity to metabolic stress in adult animals, while PPARD KO mice are embryonically lethal at mid-gestation, supporting an essential role for PPARD for the development of the organism. Cardiac-specific PPARD loss of function mice are viable, but develop severe cardiac hypertrophy at 4 weeks of age and have lower rates of FAO, thus displaying both an earlier and more severe phenotype compared to PPARA-deficient mice. Notably, overexpression of PPARA in the heart results in pathological events such as high rates of FAO and triglyceride accumulation. In contrast, overexpression of PPARD leads to increased FAO without pathological lipid storage, collectively suggesting that PPARD activation in CMs may represent an effective yet non-pathogenic approach for metabolic modulations.

In addition the isoform-specific effects observed here, one of the key findings of our work is the efficiency with which PPARD signaling is able to induce FAO in hPSC-CMs. Previous studies have established a role for PPARD signaling in mitochondrial biogenesis and regulation of lipid metabolism in mice (Kuo et al., 2013; Wang et al., 2010). In line with this, we confirm that PPARD can recapitulate the metabolic switch observed during *in vivo* maturation of the heart through enhanced FAO. These physiological changes are supported by the corresponding upregulation of the FAO machinery at multiple stages of the process. For example, CD36, the main LCFA plasma membrane transporter, is barely expressed in control hPSC-CMs, suggesting that active LCFA uptake may be limited. Upon PPARD activation however, the majority of hPSC-CMs express *CD36*, thus enabling hPSC-CMs to actively uptake LCFAs. The upregulation of *ACADVL*, which is the rate-limiting enzyme for the mitochondrial B-oxidation located in the mitochondrial membrane, further indicates an improved ability to process LCFAs. Lastly, up-regulation of *HADHA* and *HADHB*, which catalyze the last three reactions in mitochondrial B-oxidation suggests that the FAO machinery is activated from up-take all the way to the final oxidation steps.

Supplying LCFAs has been explored as an option to induce a metabolic shift in hPSC-CMs (Batho et al., 2020; Mills et al., 2017; Slaats et al., 2020; Yang et al., 2019). These studies also reported increased gene expression of components of the FAO machinery, including CD36. Our data suggest that the effects of substrate alone are indeed reproducible, but that combining LCFAs with PPARD activation leads to a stronger activation of the FAO machinery and consequently a more efficient activation of the metabolic switch. One possibility for LCFAs alone do not have a larger effect is that Carnitine palmitoyltransferase1 (CPT1) activity, encoded by *CPT1*, may be a key component in the utilization of LCFAs (Ascuitto and Ross-Ascuitto, 1996; Fisher, 1984; Warshaw and Terry, 1970). CPT1 activity changes over developmental time and acts as an indicator of when organisms begin to use LCFAs (Ascuitto and Ross-Ascuitto, 1996; Barrie and Harris, 1977; Tomec and Hoppel, 1975; Warshaw, 1970, 1972; Warshaw and Terry, 1970; Werner et al., 1983; Wittels and Bressler, 1965; Wood, 1975). For example, isolated newborn pig hearts obtain 90% of their hearts energy needs from the metabolism of circulating FAs and LCFA perfusion increases tissue levels of acyl-carnitines (Fisher, 1984; Werner et al., 1983). In many other mammals, however, CPT1 enzymatic maturation does not occur until after birth (Barrie and Harris, 1977; Tomec and Hoppel, 1975; Warshaw and Terry, 1970; Wittels and Bressler, 1965). PPARD activation in combination with LCFA supplementation strongly increases *CPT1B* expression, which is the muscle-specific CPT1 isoform, but there are no significant increases with either LCFAs or PPARD activation alone, despite strong trends in the latter condition. In the absence of increased *CPT1B* expression and activity, LCFAs alone are unlikely to extensively contribute to hPSC-CM FAO in other treatment contexts.

With the same motivation of enhancing cardiac maturation via induction of a metabolic switch, we also explored alternative metabolic modulations such as lactate selection. Nakano et al recently published an elegant study to show that glucose inhibits cardiac maturation via nucleotide biosynthesis (Nakano et al., 2017). Their data show that glucose uptake into the heart decreases during *in vivo* mouse development, starting as early as mid-gestation, and that glucose-deprivation *in vitro* in hPSC-CMs leads to enhanced cardiac maturation. Our data demonstrate enhanced FAO in the presence of glucose, and it is intriguing to speculate that a combination of glucose-deprivation with PPARD activation may lead to additive effects and further maturation of hPSC-CMs. Along similar lines and to test the effect of metabolic substrate utilization more broadly, we investigated the effect of lactate on hPSC-CM maturation. Prior to the metabolic switch to FAO, the developing heart utilizes both lactate and glucose to generate ATP in the hypoxic fetal environment (Lopaschuk and Jaswal, 2010). Furthermore, transient lactate supplementation has become a well-established protocol to efficiently enrich for hPSC-CMs and eliminate non-CM cell types based on their inability to process lactate (Ban et al., 2017; Tohyama et al., 2013). While this approach is efficient and broadly used, little is known about the mechanisms and potential effects of lactate exposure on hPSC-CMs. Here we investigated the effects of transient lactate exposure, similar to that of the lactate selection protocol for hPSC-CMs *in vitro*. We observed an unexpected reduction in the expression of components of the lactate transport and oxidation machinery 4 weeks after exposure to lactate, suggesting that hPSC-CMs display a reduced ability to uptake and process lactate as a metabolic substrate. Similar effects were described in isolated human skeletal muscle cells (Lund et al., 2018). Lactate exposure further led to downregulation of genes involved in glycolysis and the pentose phosphate pathway and resulted in a long-term reduction of LDH-A expression, outcomes which have previously been linked to maturation of hPSC-CMs (Hu et al., 2018; Nakano et al., 2017). The underlying mechanisms of why lactate exposure in hPSC-CMs leads to the changes observed here is currently unknown, and studies testing the effect of lactate during development *in vivo* are challenging. We speculate that either lactate-only, or glucose-free culture of hPSC-CM might trigger a scenario resembling starvation in cultured cells, and a subsequent activation of alternative means to generate energy, such as the oxidation of LCFAs. More work will be required to test this hypothesis, and to elucidate the detailed mechanisms of substrate modulations during hPSC-CM maturation, but the data thus far suggest that these are promising avenues to induce long term changes in hPSC-CMs, and potentially in other PSC-derived cell types.

To test whether PPARD activation and lactate selection have additive effects, we induced PPARD in lactate-treated hPSC-CMs and demonstrate that PPARD indeed induces the metabolic switch, while maintaining the gene expression changes induced upon lactate exposure. This serves to illustrate the breadth of metabolic regulation within the human heart and affirms the concept that different metabolic perturbations can have distinct yet additive effects.

An important hallmark of cardiac maturation is the increase in cardiac contractility, which *in vivo* is driven by organ and embryo growth and the related increase in circulatory demand. Several groups have illustrated that engineered tissues from hPSC-CMs can mimic this aspect *in vitro*, resulting in cells with increased contractility (Lemoine et al., 2017; Ronaldson-Bouchard et al., 2018; Tiburcy et al., 2011; Turnbull et al., 2014). Furthermore, Ulmer and colleagues have shown that enhanced contractility and maturation in engineered tissues subsequently contributes to enhanced metabolic maturation (Ulmer and Eschenhagen, 2020; Ulmer et al., 2018). We therefore hypothesized that enhanced metabolic maturation and an increase in available energy after PPARD induction should improve cardiac contractility. To test this hypothesis, we employed multiple assays which all provide parameters indicative of cardiac contractility. The combined increases in action potential duration, calcium transient duration and systolic and diastolic force in EHTs leads us to conclude that PPARD activation results not only in metabolic but also in electrophysiological and contractile maturation of hPSC-CMs, thus representing a new and easily implementable approach to efficiently induce multiple characteristics of cardiac maturation in hPSC-CMs.

In conclusion we show that PPARD activation recapitulates key developmental stages of heart development in hPSC-CMs by activating the metabolic switch from glycolysis to FAO (Table 2). PPARD signaling acts in an isoform-specific manner in hPSC-CMs with PPARD sequentially activating the gene regulatory networks underlying FAO, thus leading to increased mitochondrial content and increased LCFA up-take and processing which in turn enhances contractile maturation. Of note, PPARD activation as performed in this approach, in fully differentiated CMs, leads to stable long-term changes in hPSC-CMs, as opposed to some other maturation approaches wich assay changes earlier in the differentiation process that may be more transient. Our results thus uncover a new mechanism of *in vitro* cardiac maturation which is easily implemented and broadly applicable across many cardiac differentiation protocols. We envision that PPARD-mediated maturation will become a standalone maturation strategy with great potential to become integrated with previously established maturation strategies that will sequentially recapitulate the steps of heart development in hPSC-CMs *in vitro*. Robust *in vitro* cardiac maturation of hPSC-CMs addresses a major current roadblock and will expand the opportunities for relevant drug discovery, toxicology studies and human cardiac disease modeling.

**Table 2:** Comparison of PPARD-mediated hPSC-CM maturation and other maturation protocols. The key features of PPARD-mediated hPSC-CM maturation were compared to the changes expected during in vivo cardiac maturation and in other peer-reviewed maturation protocols. The protocols were assayed for morphology, metabolism, DNA synthesis, multinucleation, ploidy, cardiac gene expression, electrophysiology and force generation. GW0742, PPARD agonist; LCFA, Long-chain fatty acid; NS, not significant; NA, not assayed.

### Limitations of Study

In this study, we characterized hPSC-CMs 10 weeks after the start of differentiation, with PPAR modulation initiated 5 weeks after start of differentiation (3 weeks during the differentiation protocol and 2 weeks plated as monolayers). By that time, control hPSC-CMs without any PPAR activation are already relatively mature, which increases the difficulty of evaluating the cardiac maturation that we observe relative to existing peer-reviewed literature as many protocols use alternative differentiation protocols and assay specific windows of culture or differentiation time that often occur earlier during differentiation (Table 2). Overall however we believe this to equally constitute a strengths of our approach, as we observe distinct characteristics of cardiac maturation after PPARD induction even in such late stage hPSC-CMs, suggesting that these changes potentially represent a meaningful change in maturation. The overall difference in timing of the various maturation approaches (from days to weeks, some early during differentiation others weeks after start of differentiation) poses the challenge to incorporate different modifications into one protocol. Given the importance of continuously improving the still imperfect maturation state of hPSC-CMs however we propose that efforts to integrate the various discoveries and strategies will have to be designed carefully, but will prove paramount to establishing relevant and reproducible strategies in the future (Feyen et al., 2020; Funakoshi et al., 2021; Hu et al., 2018; Nakano et al., 2017; Poon et al., 2020; Tiburcy et al., 2017).

We provide here a detailed and highly mechanistic dissection of the role of substrate modulation in combination with PPAR signaling activation on cardiac maturation, and provide a new approach that results in enhanced metabolic and contractile maturation of hPSC-CMs, as well as an extensive parallel resource that we expect will facilitate further investigation of all of these mechanisms. Due to this already extensive scope, and the focus on the mechanistic aspect of the study, no disease modeling or transplantation studies were included here. It is of paramount importance however, and an exciting next task to apply the PPARD-matured hPSC-CMs to therapeutically relevant questions. CMs representing metabolically and functionally more mature cell types will likely benefit a broad range of hPSC disease models, contribute to better predictability of drug discovery and testing platforms and provide new ground for testing whether the arrhythmogenic properties of hPSC-CMs after *in vivo* transplantation can be mitigated by the generation of *in vitro* cells with enhanced maturation.

## MATERIALS AND METHODS

### Human pluripotent stem cell maintenance and differentiation

Human pluripotent stem cells (hPSCs) were maintained in E8 medium and passaged every 4 days onto matrigel-coated plates (Roche). The following hPSC lines were used in the study: MSN02-4 (human induced pluripotent stem cells generated at the Icahn School of Medicine), H9 (WA09), H7 (WA07), MEL2 and ACTN2-EGFP and MYL2-EGFP reporter lines (WTC iPSC line, GM25256) from the Allen Cell Collection, generated by the Conklin lab at the Gladstone Institutes. On Day 0 (start of differentiation) hPSCs were treated with 1 mg/ml Collagenase B (Roche) for one hour, or until cells dissociated from plates, to generate embryoid bodies (EBs). Cells were collected and centrifuged at 300 rcf for 3 min, and resuspended as small clusters of 50–100 cells in differentiation medium containing RPMI (Gibco), 2 mmol/L L-glutamine (Invitrogen), 4×104 monothioglycerol (MTG, Sigma-Aldrich), 50 µg/ml ascorbic acid (Sigma-Aldrich). Differentiation medium was supplemented with 2 ng/ml BMP4 and 3 µmol Thiazovivin (Milipore). EBs were cultured in 6 cm dishes (USA Scientific) at 37°C in 5% CO_2_, 5% O_2_, and 90% N_2_. On Day 1, the medium was changed to differentiation medium supplemented with 30 ng/ml BMP4 (R&D Systems) and 30 ng/ml Activin A (R&D Systems), 5 ng/ml bFGF (R&D Systems) and 1 µmol Thiazovivin (Milipore). On Day 3, EBs were harvested and washed once with DMEM (Gibco). Medium was changed to differentiation medium supplemented with 5 ng/ml VEGF (R&D Systems) and 5 µmol/L XAV (Stemgent). On Day 5, medium was changed to differentiation medium supplemented with 5 ng/ml VEGF (R&D Systems). After Day 8, medium was changed every 3-4 days to differentiation medium without supplements.

### Cell dissociation, plating and small molecule treatments

EBs were dissociated on day 20 of differentiation. EBs were incubated overnight with 0.6 mg/ml Collagenase Type II (Worthington) at 37°C. Dissociated cells were harvested and washed with Wash medium (DMEM, 0.1% BSA) + 1mg/ml DNase (VWR) twice and centrifuged at 300 rcf for 3 mins. Cells were resuspended in differentiation medium supplemented with 1 µmol Thiazovivin (Millipore), filtered and counted using a hemacytometer. HPSC-CMs were plated onto matrigel-coated plates at appropriate cell densities per well (6 well plate – 1,000,000 cells/well; 12 well plate – 700,000 cells/well; 48 well plate – 150,000/well; 96 well plate – 80,000 cells/well; 22×22mm coverslips for immunofluorescence – 35,000 cells/ cover slip; 22×22mm coverslips for Calcium transients – 25,000 cells/cover slip). Medium was removed the following day and replaced with differentiation medium. Medium was changed every 2 days. Two weeks after plating differentiation medium was supplemented with small molecules (GSK0660 (Sigma-Aldrich, 2.5 µM), GW0742 (Sigma-Aldrich, 5 µM), Rosiglitazone (Sigma-Aldrich, 5 µM), WY14643 (Sigma-Aldrich, 5 µM) with or without BSA-complexed long-chain fatty acids (LCFAs: FA-free BSA (Sigma-Aldrich), Palmitic Acid (Sigma-Aldrich, 12.5 µM), Oleic Acid (Sigma-Aldrich, 12.5 µM), Linoleic Acid (Sigma-Aldrich, 12.5 µM); LCFAs used in 5:4:1 ratio) for 4 weeks.

### Lactate metabolic selection

EBs were dissociated as described above and plated on Matrigel-coated 12 well plates at 700,000 cells/well in differentiation medium supplemented with 1 µmol Thiazovivin (Millipore). Medium was removed the following day and replaced with differentiation medium. After 3 days, differentiation medium was replaced with lactate medium (Stock Solution: 1 M lactate, 1 M NaHepes in distilled water; Working solution: 4mM lactate in DMEM -Glucose) for 4 days. From days 5-8, lactate medium was titrated down in the following lactate-medium : differentiation-medium ratios: Day 5: 3:1; Day 6: 1:1; Day 7: 1:3; Day 8: 0:4.

### Engineered heart tissue generation, maintenance and dissociation

EBs were dissociated by incubation with 0.6 ml/ml Collagenase Type II (Worthington) for two hours at 37°C and washed with Wash medium (DMEM, 0.1% BSA) + 1mg/ml DNase (VWR) twice and centrifuged at 300 rcf for three 3 mins. Cells were resuspended in TrypLE Express (Gibco), incubated at 37°C for three mins, washed with Wash medium (DMEM, 0.1% BSA) + 5mg/ml DNase (VWR) and centrifuged at 300 rcf for three mins. Cells were resuspended in differentiation medium supplemented with one µmol Thiazovivin (Millipore), filtered and counted using a hemacytometer. Engineered Heart Tissues (EHTs) were generated as previously described (Schaaf et al., 2014). Briefly, a cell-suspension of 1,000,000 cells, 2.5 µl Fibrinogen (Sigma-Aldrich), 3 µl Thrombin (Biopur) in 100µl differentiation medium per EHT were pipetted into molds made of 2% agarose. EHTs were then maintained in differentiation medium + Aprotinin (Sigma-Aldrich). To dissociate, EHTs were removed from posts with tweezers and dissociated overnight at 37°C in 0.6 mg/ml Collagenase Type II (Worthington). Dissociated cells were harvested and washed with Wash medium (DMEM, 0.1% BSA) + 1 mg/ml DNase (VWR) twice and centrifuged at 300 rcf for three mins. Cells were resuspended in differentiation medium supplemented with one µmol Thiazovivin, filtered and counted using a hemacytometer.

### Human Tissues

De-identified, non-anomalous second trimester human fetal heart tissue was obtained from elective terminations through the ISMMS Biorepository and Pathology Core (BRC#214).

### Immunofluorescence analysis and image processing

Tissue embedding and cryosectioning: Fetal heart tissue was fixed in 4% PFA overnight, washed with PBS, equilibrated in 30% sucrose (Sigma-Aldrich) and embedded in OCT (Electron Microscopy Services). Tissues were cut at 5 µM using a Leica Cryostat.

#### Immunofluorescence analysis

Cells plated on 22×22mm coverslips or tissue cryosections were rinsed with PBS and fixed with 4% Paraformaldehyde for 10 mins at room temperature. The edges of the coverslips were marked with a hydrophobic pen. Fixed cells were blocked with Saponin buffer (PBS, 0.5% Saponin, 0.1% BSA) for one hour at room temperature, incubated in primary antibody solution in Saponin buffer overnight at 4°C or for one hour at room temperature. Cells were rinsed three times with Saponin buffer, resuspended in secondary antibody diluted in Saponin buffer and incubated for one hour at room temperature. Cells were rinsed three times with Saponin buffer. Coverslips were mounted onto slides with nPG Antifade mounting medium (nPG stock : Glycerol : 10X PBS in the ratio 1:90:10) and edges were sealed with nail polish (VWR).

#### Primary antibody dilutions

Anti-α-Actinin (Sigma-Aldrich, 1:100), Anti-Cardiac Troponin T (Abcam, 1:400), Anti-PPARA (Santa Cruz, 1:100), Anti-PPARD (Thermo Fisher, 1:100), Anti-PPARG (Santa Cruz, 1:100), Anti-ACADVL (Abcam, 1:50), MitoTracker Deep Red dye (Thermo Fisher, 1:10,000), Anti-PMP70 AF488 (Fisher Scientific, 1:500).

#### Secondary antibody dilutions

Alexa Fluor dyes (Jackson Immunoresearch) were diluted in Saponin buffer at 1:500.

#### Image acquisition and processing

All images were acquired on a Leica DM5500 inverted microscope and processed for brightness and contrast using Fiji (Schindelin et al., 2012).

#### Image analysis

cTNT images were quantified using the MatLab-based software, MatFiber (Fomovsky and Holmes, 2010). MatFiber analyzes intensity gradients in immunofluorescent images to determine alignment (Mean Vector Length, MVL) and the circularity standard deviation (CSD). Custom FIJI and R scripts were used to determine the distance between alpha actinin+ Z-disks.

### Flow cytometry analysis and Fluorescence-Activated Cell Sorting (FACS)

#### Live flow cytometry and FACS

Dissociated cells were resuspended for 30mins on ice in differentiation medium containing antibodies (dilutions listed below). Cells were washed with differentiation medium and resuspended in differentiation medium + DAPI (1.35μg/ml, Biolegend) for flow cytometry analysis (BD LSRIIA, BD FACSCelesta) or FACS (BD FACSAria).

#### Fixed flow cytometry

Dissociated cells were fixed with 4% Paraformaldehyde for 10 mins at room temperature. Fixed cells were blocked in Saponin buffer (PBS, 0.5% Saponin, 0.1% BSA) for one hour at room temperature and resuspended in primary antibody solution in Saponin buffer overnight at 4°C. Cells were rinsed with Saponin buffer, resuspended in secondary antibody solution in Saponin buffer and incubated for one hour at room temperature. Cells were rinsed with saponin buffer and resuspended in flow buffer + DAPI for flow cytometry.

#### Antibody dilutions

PE-Cy7 anti-human CD172a/b (SIRPalpha/beta, Biolegend, 1:200), PE anti-human CD90 (Biolegend, 1:200), APC anti-human CD36 (Biolegend, 1:100), MitoTracker Deep Red dye (Thermo Fisher, 1:10,000; binds mitochondria in a delta-psi independent manner), Anti-Cardiac Troponin T (Abcam, 1:400), PMP70 AF488 (Fisher Scientific, 1:500).

### cDNA generation and real-time qPCR

RNA was isolated from cells/tissues using the Quick-RNA kit (Zymo Research). cDNA was generated using reverse transcription with the Quanta qScript kit (Quanta bio). Quantitative PCR was carried out on an Applied Biosystems Step One Plus using ABI SYBR Green reagents. Expression levels were normalized to TATA-binding protein (TBP) and genomic DNA was used for quantification of absolute expression levels, as previously described (Dubois et al., 2011).

### RNA sequencing

RNA was extracted from FACS-isolated SIRPA+CD90-hPSC-CMs using the Quick RNA Micro kit (Zymo Research). 500 ng of RNA per sample was used for Automated RiboZero Gold library prep and sequenced using a S2 100 cycle flow cell. FASTQ files were aligned with kallisto to a reference transcriptome model generated from the human genome hg38 using default parameters (Bray et al., 2016). The average number of reads successfully pseudoaligned across samples was 16.46 million, with a standard deviation of 3.76 million. DESeq2 was used to normalize read counts and determine differentially expressed genes with p<0.05. GAGE was used to identify GO and KEGG pathways that were upregulated or downregulated with p<0.05. Custom R scripts were used to generate heatmaps. Published code was adapted to generate UpsetR plots, KEGG comparisons and Chord Plots (Lex et al., 2014).

### ATAC Sequencing

ATACseq was performed as described in (Buenrostro et al., 2015). Briefly, cells were FACS-isolated and split into aliquots of 50,000 cells from each sample. After washing with PBS, samples were resuspended in cold lysis buffer (10 mM Tris-HCl pH 7, 10 mM NaCl, 3 mM MgCl2, 0.01% Igepal CA-630) and spun down at 500 x g for 10 min at 4°C. Cells were incubated with 25 μL TD buffer, 2.5 μL Tn5 transposase (Illumina; Cat No: #20034198) and 22.5 μL nuclease-free H20 at 37°C for 30 min with gentle agitation. After purification (Qiagen; Cat No: #28004), transposed DNA fragments were amplified with custom PCR primers (Buenrostro et al., 2013) for a total of 9 cycles. The final PCR cycle number needed to minimally amplify libraries was determined by a separate qPCR reaction as described in Buenrostro et al., 2015 to minimize GC and size bias. Purified libraries were sequenced on an Illumina Novaseq 6000. Adapters were trimmed from the raw fastq data with cutadapt, and the raw data was aligned to hg38 with bwa mem. PCR duplicates were removed with Picard, in addition to reads mapping to mitochondrial sequences or known problematic genomic regions in the ENCODE blacklist. Macs2 was used to call peaks with the bampe parameter. To reduce false positives, peaks were used when overlapping with consensus regions mapped in at least two samples. Featurecounts was used to count reads associated with known promoter regions or candidate regulatory elements listed in the SCREEN (Search Candidate cis-Regulatory Elements by ENCODE) database. DESeq2 was used to identify differential peaks with p<0.05, and peaks were annotated with the closest genes.

### Seahorse analysis

Dissociated cells were plated at 25,000 cells per well in a 96-well microplate (Agilent). One week after plating, an XF Cell Mito Stress Test (Agilent, Oligomycin 1 µM, FCCP 2 µM, Rotenone and Antimycin A 0.5 µM) was performed using an XFe96 Analyzer. Cell numbers were normalized using Methylene Blue staining and normalized data were analyzed using Seahorse Wave Desktop Software.

### Fatty acid oxidation flux assay

The FAO flux assay quantifies the rate of [9,10-3H] Palmitate (PerkinElmer, Waltham, MA) oxidation. [9,10-3H] Palmitate is emulsified in Kreb/BSA solution (4.5 mg/ml) to a final concentration of 23 μM (Master Mix) overnight at 370C under continuous agitation. FACS-isolated SIRPA+CD90-hPSC-CMs were seeded in a 96-well plate (80,000cell/well) and were incubated for 4h with 200 μl Master Mix. The 3H2O released into the cell medium was quantified. Etomoxir (2[6(4-chlorophenoxy) hexyl]oxirane-2-carboxylate, 10 µM) an irreversible inhibitor of carnitine palmitoyltransferase-1 (CPT-1a) was used to document the mitochondrial-dependent β-oxidation of palmitate. The flux is expressed in pmol/h/mg protein.

### Multi-electrode array analysis

Dissociated cells were plated at 60,000 cells per well in a CytoView 48-well multielectrode array (MEA) plate (Axion Biosystems). One week after plating recordings were taken using an Axion Integrated Studio (AxIS) on the Axion Maestro. Cardiac Beat Detector measurement parameters were set at 300 μV detection threshold, 250 ms min. beat period, 5s max. beat period, polynomial regression FPD detection method, 70 ms post-spike detection holdoff, 50 ms pre-spike detection holdoff, 1s match post search duration with limit FPD search, and 10 running average beat count used for FPD detection.

### Intracellular calcium transient (CaT) and action potential (AP) analysis

FACS-isolated SIRPA+CD90-hPSC-CMs were plated at 30,000 cells/coverslip (Sarstedt). 7 days post-plating AP and CaT measurements were performed using voltage and Ca2+ sensitive dyes. Briefly, coverslips were incubated in 1X Tyrodes’ buffer (140 mM NaCl, 5.4 mM KCl, 10mM HEPES, 1 mM NaH2PO4, 1 mM MgCl2, 10 mM Glucose and 1.8 mM CaCl2, calibrated to pH=7.4) containing 10 mM Calbryte-630 AM (AAT Bioquest) and Fluovolt dyes (Invitrogen – dilution 1:1000 Fluovolt dye, 1:100 Powerload concentrate) in order to get paired AP and CaT recordings from a single coverslip. Coverslips were incubated in dye solution for 30 minutes at 37°C. Coverslips were then transferred to custom cassette insert and incubated in 1X Tyrode’s buffer with 10 mM blebbistatin for 5 minutes, then loaded onto microscope for measurement. Tyrodes’ buffer was perfused throughout measurement to maintain cells at 37°C. Cells were paced at 0.5 Hz using a MyoPacer Field Stimulator. 10-15 cells were recorded per coverslip across three biological replicates, resulting in a total of 30-45 cells per condition. Recordings of fluorescence flux in line scan were taken on an LSM5 exciter microscope using Zeiss ZEN 2009 software. Image files were analyzed using custom MATLAB scripts to extract features from individual waveforms for quantification.

### Engineered heart tissue deflection analysis

Engineered Heart Tissues (EHTs) were generated using established protocols and described above (Schaaf et al., 2014). 14 days post-fabrication, contractile function was measured every week for 5 weeks (from week 1-4 of treatment). At the day of measurement, pacing electrodes were positioned along each array of tissues and the plate was put into the incubator to equilibrate (Mannhardt et al., 2017). After 30 minutes, the plate was placed on a stage equipped with a right angle mirror reflecting the image of the EHT posts to a dissecting microscope and high-speed camera (Turnbull et al., 2018). LabVIEW software was used to acquire real-time data of post deflection, by applying a beam-bending equation from elasticity theory as previously described (Serrao et al., 2012). The data were analyzed with a custom MATLAB script to calculate twitch parameters including developed force (DF; calculated from the post deflection with each twitch), maximum contraction rate (+dF/dt) and maximum relaxation rate (-dF/dt), as previously described (Turnbull et al., 2014). Functional data were first recorded under spontaneous conditions, including measuring spontaneous beating rate. For paced measurements the electrodes were connected to a S88X Grass Stimulator (Astro-Med, Inc., West Warwick, RI) and data were recorded under electrical stimulation, at rates from 0.25 Hz to 3.0Hz with 0.25Hz increments. At the end of each day of recording, the electrodes were removed and medium was replaced with fresh medium.

### Transmission electron microscopy

Dissociated cells were plated onto 22×22mm coverslips such that they covered the majority of the coverslip surface. The cells were fixed with 2% paraformaldehyde and 2% glutaraldehyde/PBS, pH 7.2 for a minimum of one week at 4 degrees C. Sections were rinsed in 0.1 M sodium cacodyl-ate buffer (EMS), fixed with 1% osmium tetroxide followed with 2% uranyl acetate, dehydrated through ascending ethanol series (beginning with 25% up to 100%) and infiltrated with Embed 812, an epon resin kit (EMS). Beem capsules (EMS) were placed on top of the cells, filled with resin, and heat polymerized at 60 degrees C for 72 hrs. Post polymerization, capsules were heated to 60 degrees C for 3.5 minutes and snapped from the substrate to dislodge the cells. Semithin sections (0.5 and 1 µm) were obtained using a Leica UC7 ultramicrotome, counterstained with 1% Toluidine Blue, cover slipped and viewed under a light microscope to identify successful dislodging of cells. Ultra-thin sections (80nms) were collected on nickel 300 mesh grids (EMS) using a Coat-Quick adhesive pen (EMS). Sections were counter-stained with 1% uranyl acetate followed with lead citrate, and imaged on an Hitachi 7000 electron microscope (Hitachi High-Technologies, Tokyo, Japan) using an advantage CCD camera (Advanced Microscopy Techniques, Danvers, MA). Images were adjusted for brightness and contrast using Adobe Photoshop CS4 11.0.1.

### Statistical Analyses

Data are presented as mean ± standard deviation. Statistical significance between two groups was determined using two-sided unpaired or paired Student’s t-tests. One-way ANOVAs were used when testing for relationships between 3 or more groups with appropriate post-hoc testing. The statistical tests performed are included in the figure legends. Statistical tests were performed using GraphPad Prism 6.

## Data availability

The authors declare that all data supporting the findings of this study are available within the article and its supplementary information files, or from the corresponding author upon request. The RNA sequencing data are deposited in the NCBI GEO database. The accession code is GSE160987. The ATAC sequencing data are deposited in the NCBI GEO database. The accession code is GSE178984.

### ACKNOWLEDGMENTS

We thank the ISMMS Flow Cytometry Core (Christopher Bare, Xuqiang Qiao), the ISMMS Microscopy Core (William Jansen, Allison Sowa, Esperanza Agullo Pascual, Shilpa Dilipkumar, Deanna Benson), the ISMMS Biorepository and Pathology Core (Michael Donovan, Olha Fedorshyn, Anastasiya Dzuhun), the ISMMS Stem Cell Shared Resource facilities and the NYU Genome Technology Center (Adriana Heguy, Sitharam Ramaswami) for their technical assistance. Jinqi Gong and Jaehee Shim (Sobie Lab, ISMMS) provided assistance with calcium transient measurements and analysis. Irene C. Turnbull is ethically opposed to research involving human embryonic stem cells and tissues derived from elective abortions; Dr. Turnbull’s contribution to this study included performing the functional measurements and data analysis of human engineered tissues, and instruction on use of MatFiber to quantify myofibril organization. We thank her for her invaluable contributions to this project. The LINCS Consortium [NIH/NHLBI 5U54HG008098-02 to Ravi Iyengar] provided the MSN02-4 hiPSC line.

## AUTHOR CONTRIBUTIONS

NMW and NCD designed and performed experiments and analyzed data. DS, DT and AM analyzed RNAseq and ATACseq data. DMG, BS, PD, EM, SR, RD, MS and JM performed experiments and data analysis. RS, KB and SI designed and performed the scRNAseq experiments. DF, HI and SH provided advice on gene editing and metabolism experiments. AJ assisted in obtaining fetal heart tissue. AH and TE shared expertise and materials for generating engineered heart tissues from hPSC-CMs. NMW and NCD wrote the manuscript with input from all authors.

## SUPPLEMENTARY FIGURES

**Figure S1:**
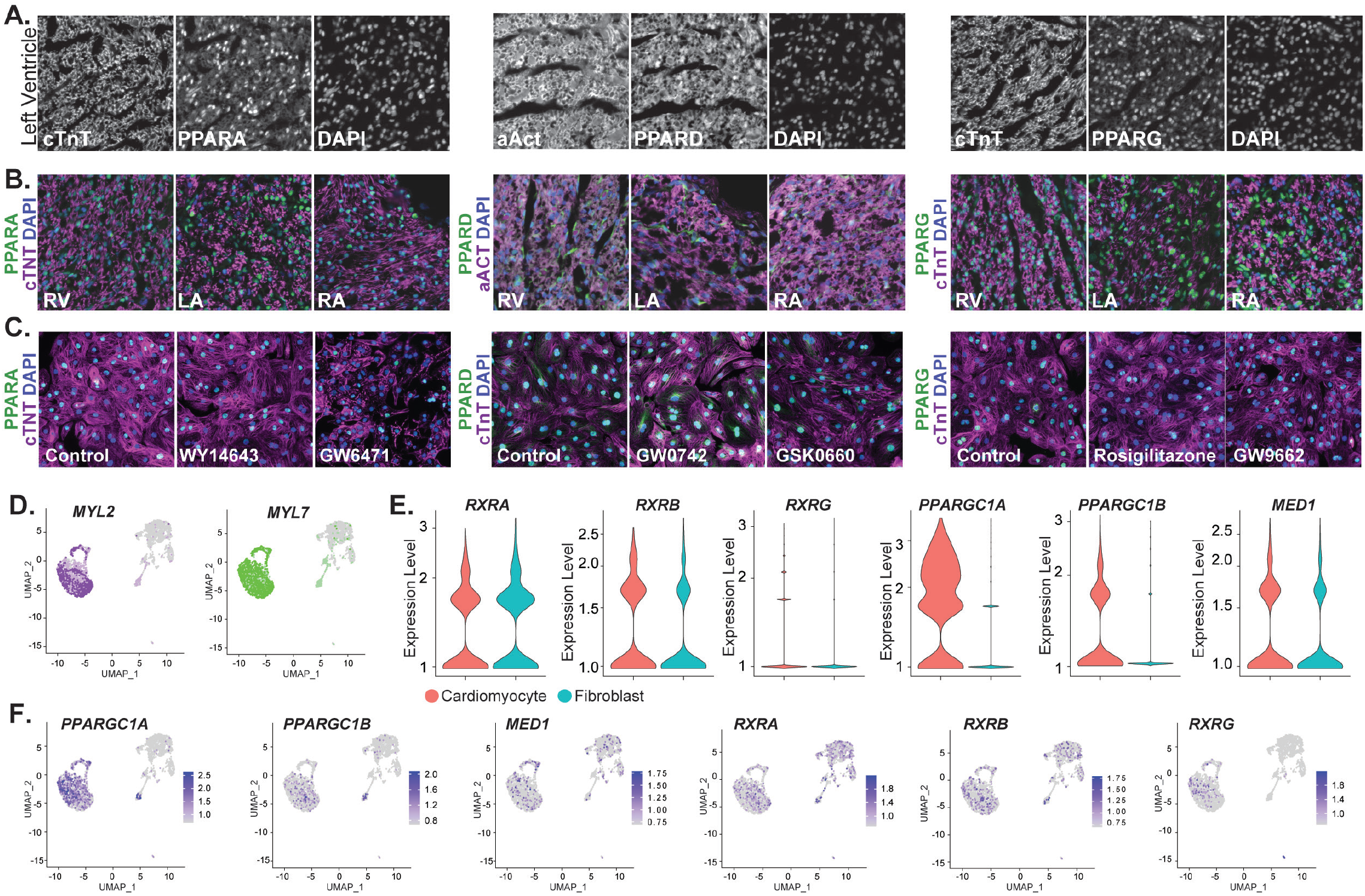
Expression of the PPAR signaling machinery in hPSC-CMs and fetal heart tissues. (**A**) IF analysis for cardiac Troponin T (cTnT) or alpha-actinin (aAct) in combination with PPARA, PPARD and PPARG on week 16 human heart left ventricle sections. DAPI is used to visualize cell nuclei. Scale bar = 25μm. (**B**) IF analysis for cTnT or alpha-actinin in combination with PPARA, PPARD and PPARG on week 16 human heart right ventricle (RV), left atrium (LA) and right atrium (RA) sections. DAPI is used to visualize cell nuclei. Scale bar = 25μm. (**C**) IF analysis for cardiac cTnT in combination with PPARA, PPARD and PPARG on 30-day-old FACS-sorted hPSC-CMs. DAPI is used to visualize cell nuclei. Scale bar = 75μm. (**D**) ScRNAseq UMAP clustering analysis showing expression of *MYL2* (purple) and *MYL7* (green) in hPSC-CMs. (**E**) Expression of *RXRA, RXRB* and *RXRG* in hPSC-CMs and hPSC-Fibroblasts. (F) ScRNAseq UMAP showing expression of *PPARGC1A, PPARGC1B, MED1, RXRA, RXRB* and *RXRG* in hPSC-CMs.

**Figure S2:**
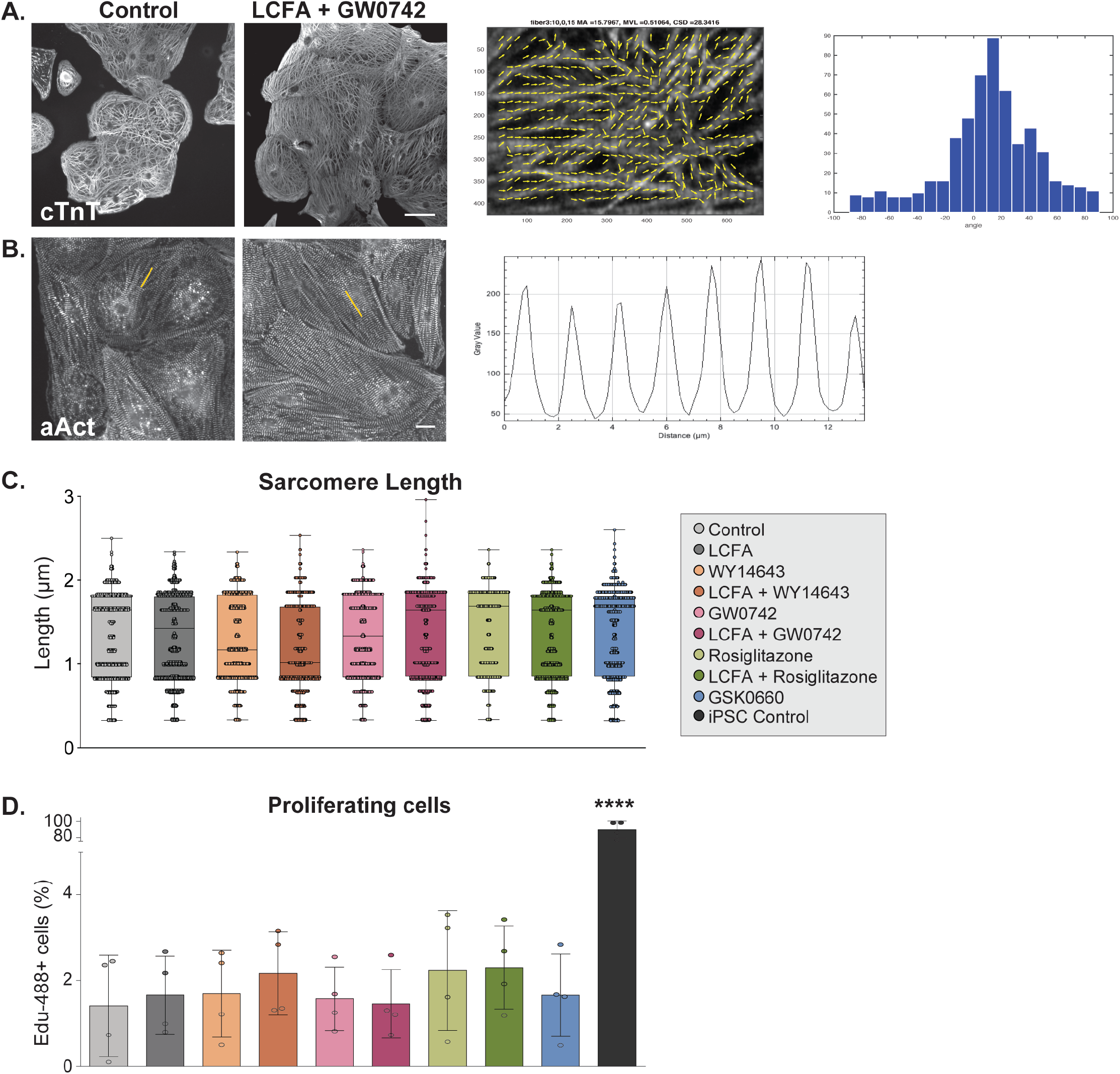
PPARD signaling modulation does not affect sarcomere lengths or proliferation rate of hPSC-CMs. (**A**) IF analysis for cTnT on hPSC-CMs 4 weeks after continuous PPARD modulations (left). Images were processed by MatFiber to determine Mean Vector Length (MVL) and Circularity Standard Deviation (CSD). Representative plot of the distribution of vectors between -180o and 180o (middle). Representative image of grey value intensity along an Region of Interest (ROI)(right). (**B**) IF analysis on hPSC-CMs 4 weeks after PPAR modulations for aAct. Representative images generated from aAct IF in ImageJ to determine distance between sarcomeres (left). Quantification of sarcomere length in hPSC-CMs 4 weeks after PPAR modulations (right). Yellow line indicates example of myofibril chosen to measure sarcomere lengths. (**C**) Quantification of sarcomere lengths in control and PPAR-modulated hPSC-CMs 4 weeks after treatment (2000 < n < 4000). (**D**) Differentiation cultures were incubated for 24 hours with EdU at the end of 4 weeks of PPAR modulation. Edu incorporation was determined in cardiac cTnT+ hPSC-CMs via flow cytometry analysis (n=4). Data represented as mean + SD. Statistics: One-way ANOVA and Tukey test for multiple comparisons relative to the control condition (far left)(*p<.05; **p<.01); ***p<.001; **** p<.0001).

**Figure S3:**
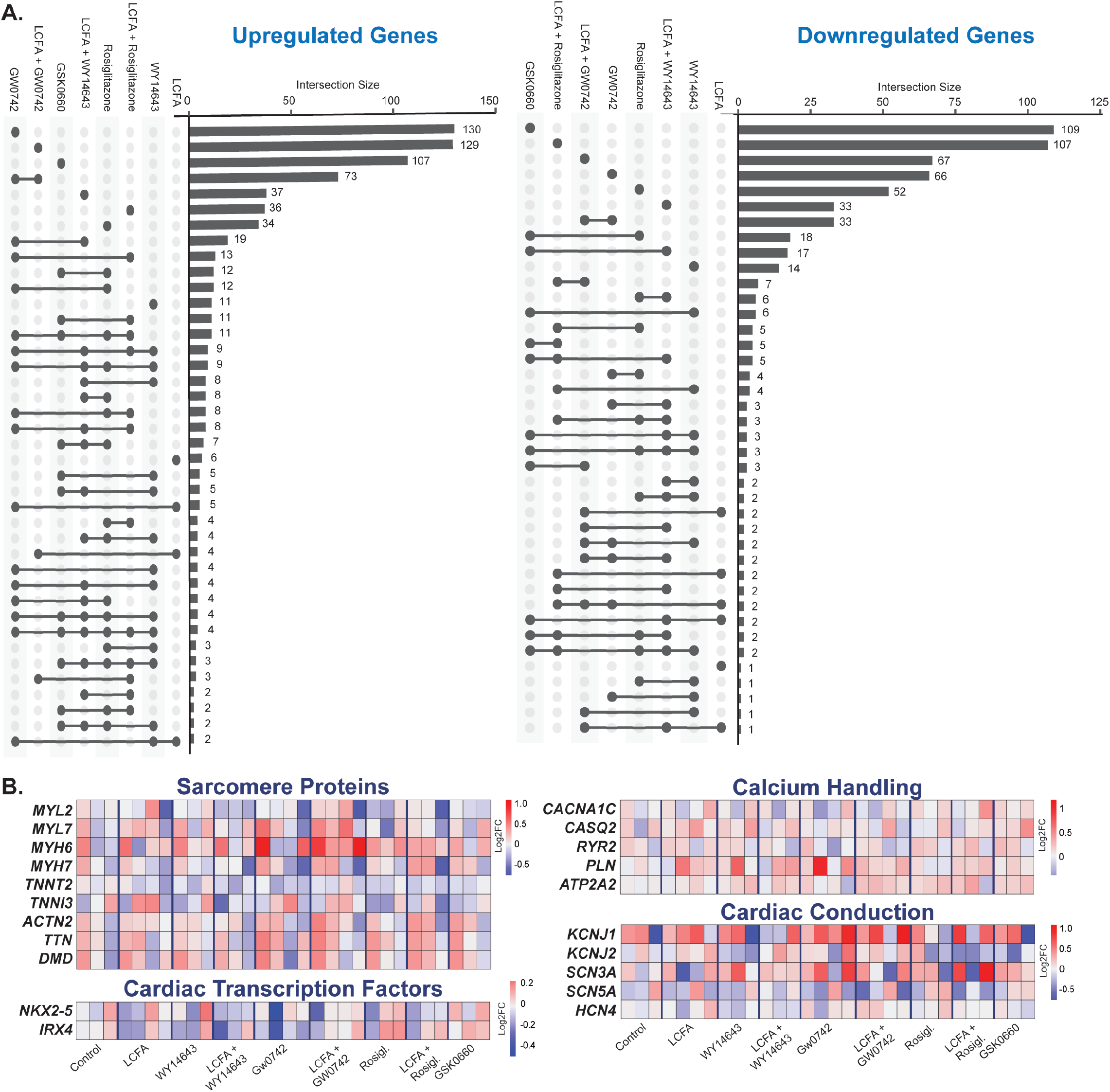
Changes in gene expression after PPAR induction/inhibition are isoform-dependent. (**A**) UpsetR plots summarizing the number of differentially expressed genes (compared to control hPSC-CMs) that are shared between different culture modulations (left: upregulated, right: downregulated). Statistics: DESeq2 was used to normalize read counts and determine differentially expressed genes with p<0.05. (**B**) Gene expression analysis in hPSC-CMs 4 weeks after continuous PPARD modifications (RNAseq, n=3). Select candidates are shown for sarcomere structure (*MYL2, MYL7, MYH6, MYH7, TNNT2, TNNI3, ACTN2, TTN, DMD*), calcium handling (*CACNA1C, CASQ2, RYR2, PLN, ATP2A2*), cardiac conduction (*KCNJ1, KCNJ2, SNC3A, SCN5A, HCN4*) and cardiac transcription factors (*NKX2-5, IRX4*). Statistics: DESeq2 was used to normalize read counts and determine differentially expressed genes with p<0.05 relative to the control condition (far left).

**Figure S4:**
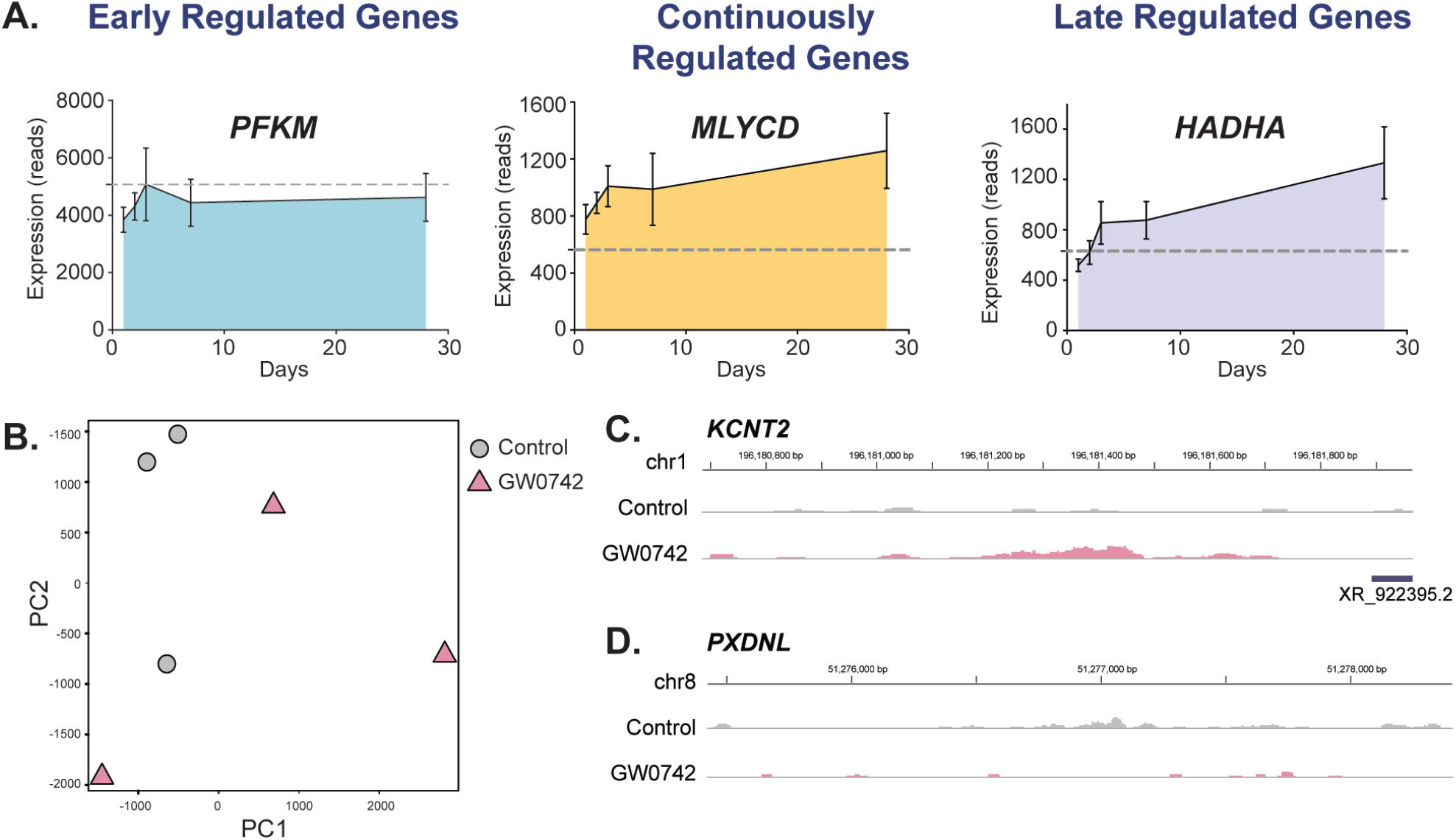
PPARD activation induces time-dependent transcriptional changes and alters chromatin accessibility. (**A**) Gene expression analysis of candidates that display a dependency of GW0742 treatment duration: Early regulation: PFKM, Continued regulation: MLYCD, Late regulation: HADHA. Control expression levels are represented as a gray dashed line (n=3). (**B**) Principal Component Analysis (PCA) of ATACseq data from control and GW0742-treated hPSC-CMs after 28 days of treatment, showing the separation of the two populations along the PC1 vs. PC2 axes (n=3). (**C**) ATACseq tracks of *KCTN2* and *PXDNL* displaying differential chromatin accessibility in promoter and enhancer associated regions in control and GW0742-treated hPSC-CMs. Data represented as mean + SD. Statistics: DESeq2 was used to normalize read counts and determine differentially expressed genes with p<0.05 relative to the control condition (far left).

**Figure S5:**
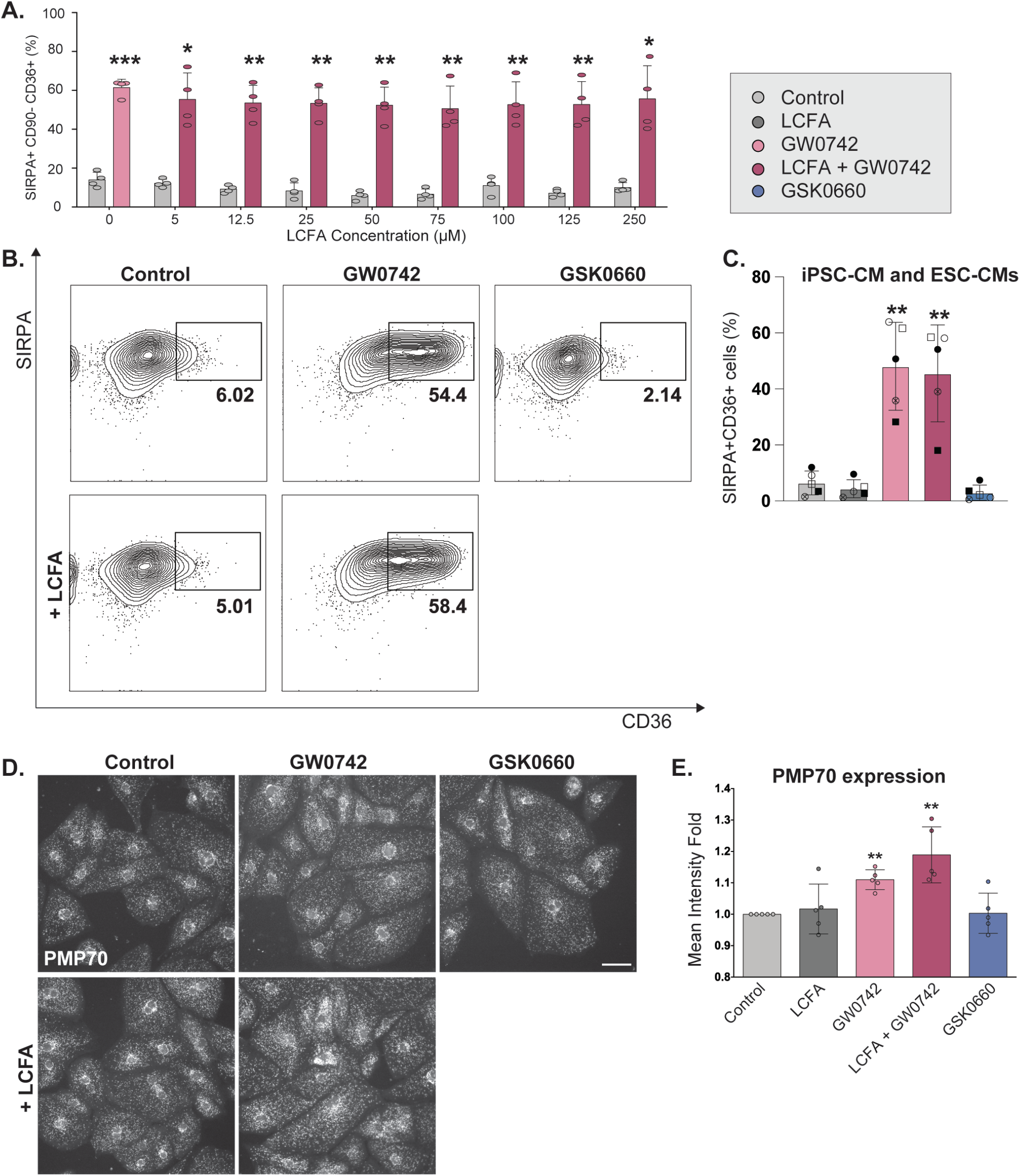
PPARD activation increases CD36 cell surface expression across multiple hPSC lines and LCFA concentrations, and increases peroxisome content. (**A**) Quantification of flow cytometry analysis of hPSC-CMs 4 weeks after continuous PPAR modulations supplemented with BSA-complexed LCFAs from 0-250μM. SIRPA: cardiomyocyte marker, CD36: Fatty acid transporter (n=4 biological replicates). (**B**) Representative flow cytometry analysis of hPSC-CMs derived from H9 4 weeks after continuous PPARD modulations. (**C**) Quantification of flow cytometry analysis (SIRPA+CD36+) of GW0742-treated hPSC-CMs (4 weeks) from different hiPSC and hESC lines. Each data point represents a different biological replicate (n=5, 3 hESC (circles, H9), 2 iPSC (squares, GM25256). (**D**) IF analysis for PMP70, a peroxisome-specific protein, in hPSC-CMs 4 weeks after continuous PPARD modulations. (**E**) Mean fluorescence intensity for PMP70 was quantified through flow cytometry in hPSC-CMs 4 weeks after continuous PPARD modulations (n=4). Scale bars: 50µM. Data represented as mean + SD. Statistics: Student’s t-tests relative to the control condition (far left) (*p<.05; **p<.01; ***p<.001; **** p<.0001).

**Figure S6:**
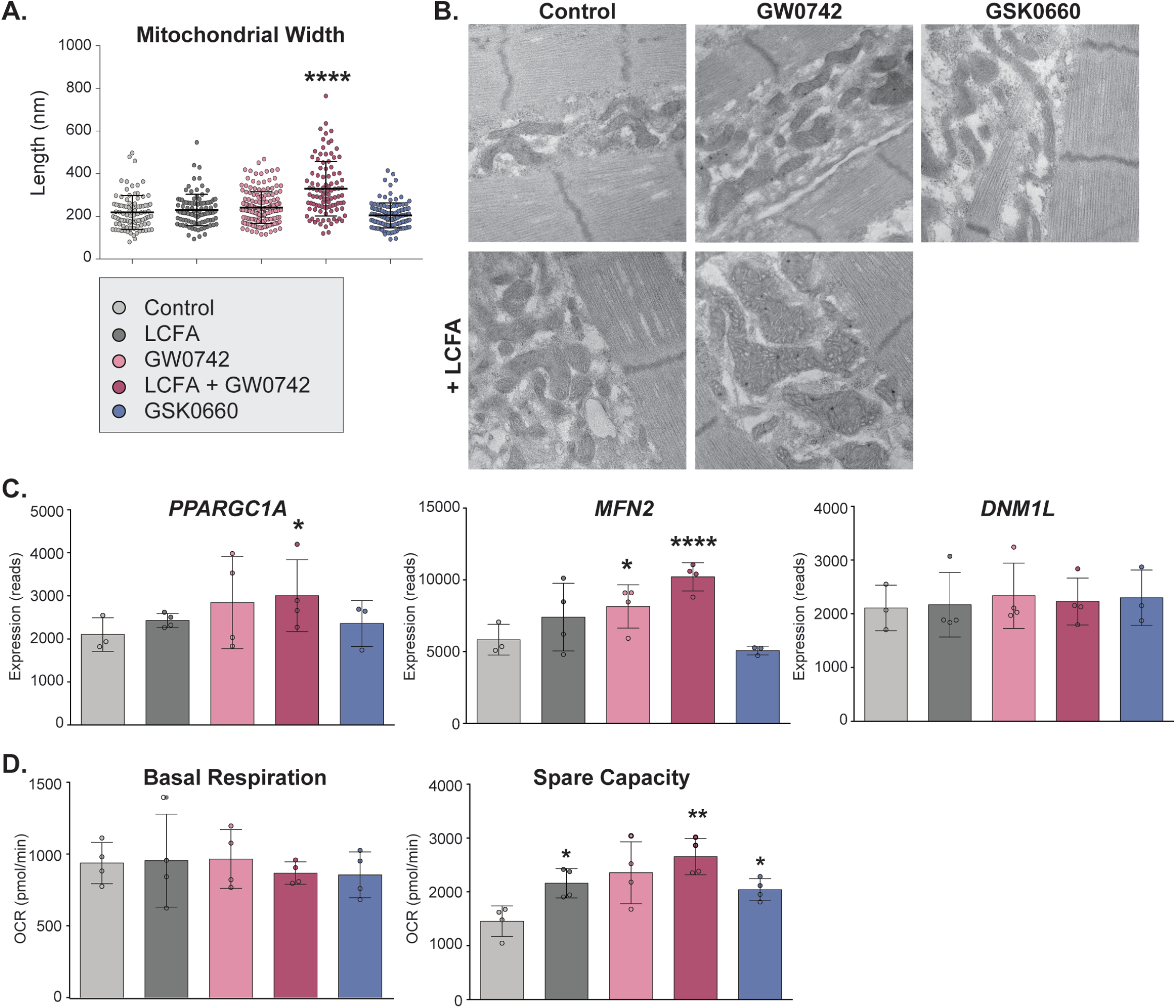
PPARD signaling results in enhanced mitochondrial ultra-structure and up-regulation of candidates involved in mitochondrial biogenesis and fusion. (**A**) Quantification of mitochondrial width from images in Figure 5 (n>99/condition). Data represented as mean + SD. Statistics: One-way ANOVA and Tukey test for multiple comparisons (*p<.05; **p<.01); ***p<.001; **** p<.0001). (**B**) Transmission electron microscopy imaging of hPSC-CMs 4 weeks after continuous PPARD modifications to illustrate mitochondrial cristae formation. (**C**) Gene expression analysis of hPSC-CMs 4 weeks after PPARD modulations (RNAseq, n=3) for candidates important for mitochondrial biogenesis including *PPARGC1A, MFN2*, and *DNM1L*, involved in mitochondrial biogenesis, fusion and fission respectively. Data represented as mean + SD. Statistics: DESeq2 was used to normalize read counts and determine differentially expressed genes with p<0.05. (**D**) Oxygen consumption rate (OCR) measured with a Seahorse analyzer in hPSC-CMs 4 weeks after PPARD modulations. Basal respiration (left) and respiratory spare capacity (right) are shown (n=4; 5-8 wells/condition per biological replicate). Data represented as mean + SD. Statistics: Student’s t-tests relative to the control condition (far left) (*p<.05; **p<.01; ***p<.001; **** p<.0001).

**Figure S7:**
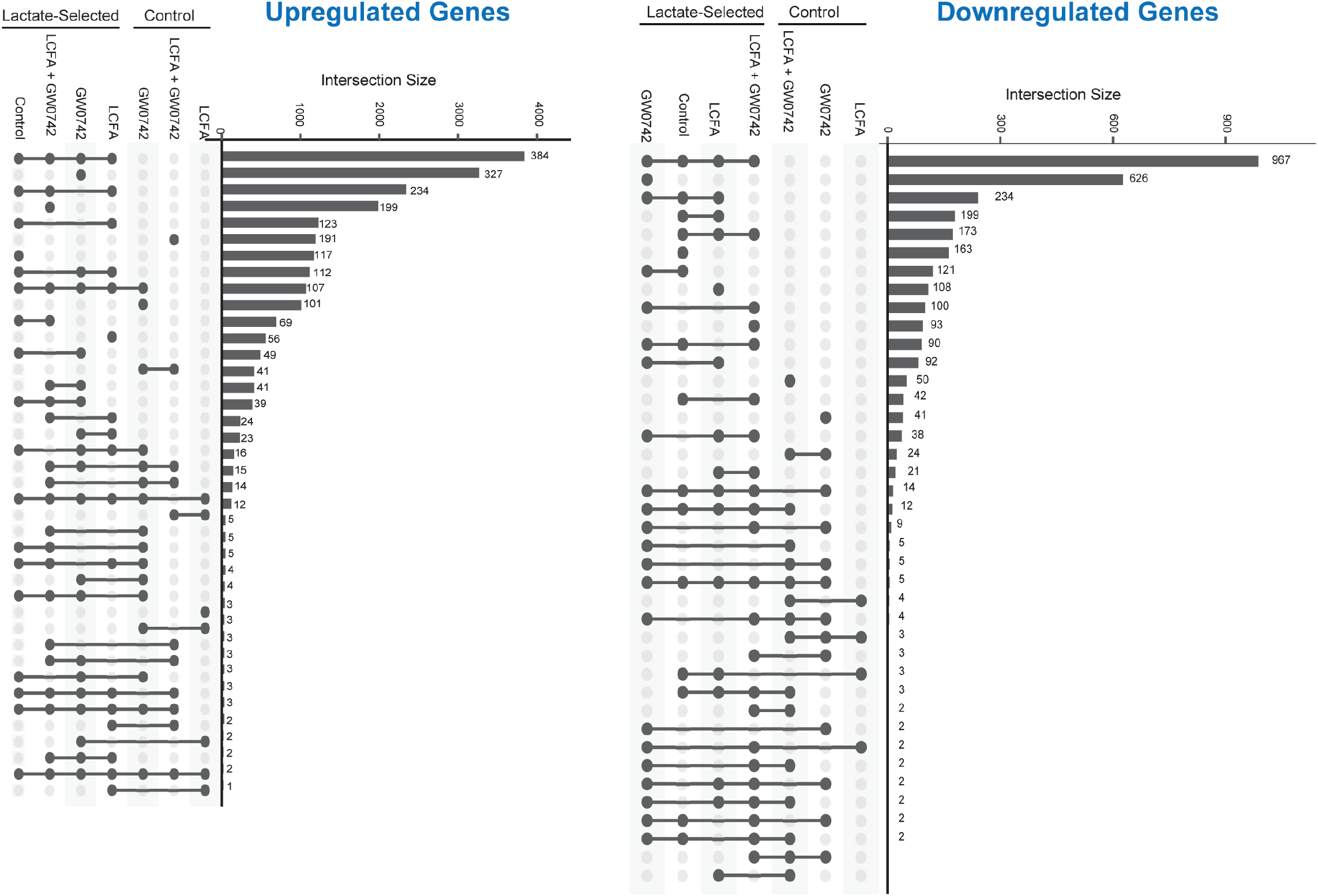
Differential gene expression analysis in hPSC-CMs after lactate selection in combination with PPARD modulation. UpsetR plots summarizing the number of differentially expressed genes (compared to control hPSC-CMs) after lactate selection and in combination with different PPAR modulations (left: upregulated, right: downregulated). Statistics: DESeq2 was used to normalize read counts and determine differentially expressed genes with p<0.05.

**Supplemental File 1: Gene lists of early, maintained and late genes after PPARD signaling induction**. Log2FC differences relative to control after 1, 2, 3, 7 and 28 days of PPARD activation.

**Supplemental File 2: Lists of up-regulated and down-regulated genes after transient PPARD signaling induction**. Gene list of genes differentially regulated after 7d, 7d-pulse or 28 days of PPARD Ag treatment relative to the 28-day control condition.

Supplemental File 3: EHT morphology after PPARD signaling activation/inhibition. Representative videos hPSC-CM EHTs 4 weeks after PPAR modulations.

