## Supplementary material for "PPARdelta signaling activation improves metabolic and contractile maturation of human pluripotent stem cell-derived cardiomyocytes": Table1

| **KEGG Term** | **Description** | **Early** | | | **Continued** | | | **Late** | |
| --- | --- | --- | --- | --- | --- | --- | --- | --- | --- |
|  |  | **No.** | **Description** | **No.** | | **Description** | **No.** | | **Description** |
| **Upregulated** | |  |  | |  |  | |  |  |
| hsa03320 | PPAR signaling pathway | 2 | ACOX3 SCP2 | | 1 | CPT1A | | 6 | DBI ACSL1 ANGPTL4 SCD HMGCS2 CD36 |
| hsa04110 | Cell cycle | 17 | MCM7 E2F3 CDC7 CHEK1 PKMYT1 SMC1A CDC45 CDC26 CDKN1C E2F5 TFDP1 CDKN1C YWHAQ ORC1 E2F2 CDC6 PRKDC YWHAZ E2F2 | | 0 |  | | 0 |  |
| hsa03030 | DNA replication | 8 | MCM7 POLA2 POLE PRIM1 LIG1 RPA3 FEN1 PRIM2 | | 0 |  | | 0 |  |
| hsa03440 | Homologous recombination | 5 | BRCA2 MRE11 RAD51 RPA3 EME1 | | 0 |  | | 0 |  |
| hsa03430 | Mismatch repair | 4 | MSH3 EXO1 LIG1 RPA3 | | 0 |  | | 0 |  |
| hsa00240 | Pyrimidine metabolism | 11 | POLA2 DUT TYMS POLE DCK CTPS1 RRM1 PRIM1 TK1 DTYMK PRIM2 | | 1 | AK3 | | 0 |  |
| hsa04114 | Oocyte meiosis | 8 | REC8 PKMYT1 SMC1A MAPK1 CDC26 YWHAQ YWHAZ IGF1R REC8 | | 0 |  | | 0 |  |
| hsa00190 | Oxidative phosphorylation | 2 | NDUFB5 SDHD | | 0 |  | | 0 |  |
| hsa04010 | MAPK signaling pathway | 6 | NRAS FGFR2 MAPK1 RRAS2 FGF1 MAPK8 | | 0 |  | | 1 | PLA2G12A |
| hsa04141 | Protein processing in endoplasmic reticulum | 4 | UBE2E1 UBE2E3 UBE2D3 MAPK8 | | 0 |  | | 0 |  |
| hsa04380 | Osteoclast differentiation | 3 | MAPK1 FYN MAPK8 | | 0 |  | | 0 |  |
| hsa04612 | Antigen processing and presentation | 1 | RFX5 | | 0 |  | | 0 |  |
| hsa04145 | Phagosome | 3 | TUBB TUBB TUBB TUBB TUBA1B TUBB TUBB TUBB STX7 TUBB | | 0 |  | | 1 | CD36 |
| hsa04012 | ErbB signaling pathway | 4 | NRAS MAPK1 MAPK8 PAK3 | | 0 |  | | 0 |  |
| hsa04620 | Toll-like receptor signaling pathway | 2 | MAPK1 MAPK8 | | 0 |  | | 0 |  |
| hsa04920 | Adipocytokine signaling pathway | 1 | MAPK8 | | 1 | CPT1A | | 2 | ACSL1 CD36 |
| hsa00500 | Starch and sucrose metabolism | 0 |  | | 0 |  | | 1 | UGP2 |
| hsa04662 | B cell receptor signaling pathway | 2 | NRAS MAPK1 | | 0 |  | | 0 |  |
| hsa04650 | Natural killer cell mediated cytotoxicity | 3 | NRAS MAPK1 FYN | | 0 |  | | 0 |  |
| hsa00052 | Galactose metabolism | 0 |  | | 0 |  | | 1 | UGP2 |
| hsa04260 | Cardiac muscle contraction | 1 | TPM4 | | 0 |  | | 0 |  |
| hsa00520 | Amino sugar and nucleotide sugar metabolism | 1 | GNPDA2 | | 0 |  | | 1 | UGP2 |
| hsa04350 | TGF-beta signaling pathway | 11 | SMAD1 SMAD7 ZFYVE16 MAPK1 DCN E2F5 TFDP1 FST ID3 AMHR2 SMAD6 ID3 | | 0 |  | | 0 |  |
| hsa04144 | Endocytosis | 8 | SMAD7 ZFYVE16 FGFR2 MVB12B CHMP2B KDR SMAD6 IGF1R | | 0 |  | | 0 |  |
| hsa04660 | T cell receptor signaling pathway | 4 | NRAS MAPK1 FYN PAK3 | | 0 |  | | 0 |  |
| hsa04972 | Pancreatic secretion | 1 | TRPC1 | | 0 |  | | 1 | PLA2G12A |
| **Downregulated** | |  |  | |  |  | |  |  |
| hsa03320 | PPAR signaling pathway | 2 | PPARA ACOX2 | | 0 |  | | 0 |  |
| hsa00260 | Glycine, serine and threonine metabolism | 2 | AMT ALAS1 | | 2 | CBS GCAT | | 0 |  |
| hsa04110 | Cell cycle | 5 | EP300 CCND1 YWHAE WEE1 YWHAE CDKN2D | | 0 |  | | 0 |  |
| hsa03030 | DNA replication | 1 | POLE3 | | 0 |  | | 0 |  |
| hsa00240 | Pyrimidine metabolism | 3 | PNP POLR3D POLE3 | | 0 |  | | 0 |  |
| hsa04114 | Oocyte meiosis | 5 | ADCY5 YWHAE PPP2R5D YWHAE PPP2CB ADCY6 | | 1 | RPS6KA2 | | 0 |  |
| hsa00010 | Glycolysis / Gluconeogenesis | 4 | ALDH3B1 PFKM ENO1 PFKL | | 0 |  | | 0 |  |
| hsa00190 | Oxidative phosphorylation | 1 | ATP6V0C | | 0 |  | | 0 |  |
| hsa04010 | MAPK signaling pathway | 15 | CRKL DDIT3 CRK PAK1 DUSP6 DUSP8 DUSP5 DUSP2 DAXX DAXX DAXX ATF4 DUSP8 FGF5 DUSP8 DUSP10 FGF18 DUSP1 DAXX DAXX DUSP7 DAXX | | 1 | RPS6KA2 | | 1 | FGFR2 |
| hsa04141 | Protein processing in endoplasmic reticulum | 10 | DDIT3 DERL3 AMFR EIF2AK4 EIF2AK3 DERL3 CAPN2 ATF4 EDEM1 DDOST PPP1R15A | | 1 | SYVN1 | | 0 |  |
| hsa04380 | Osteoclast differentiation | 5 | SQSTM1 PIK3CD TNFRSF11B TEC FOSL2 | | 0 |  | | 1 | CYBA |
| hsa04976 | Bile secretion | 3 | ADCY5 ATP1B2 ADCY6 | | 1 | ATP1A2 | | 0 |  |
| hsa00051 | Fructose and mannose metabolism | 4 | PMM1 PFKFB3 PFKM PFKL | | 0 |  | | 0 |  |
| hsa04610 | Complement and coagulation cascades | 2 | C1R C1S C1R | | 0 |  | | 0 |  |
| hsa04145 | Phagosome | 7 | TFRC C1R DYNC1H1 ATP6V0C TUBA1C TUBB4B PIKFYVE C1R | | 0 |  | | 1 | CYBA |
| hsa04012 | ErbB signaling pathway | 8 | CRKL CRK PAK1 ERBB3 PIK3CD PAK4 EREG STAT5A | | 0 |  | | 0 |  |
| hsa04620 | Toll-like receptor signaling pathway | 1 | PIK3CD | | 0 |  | | 0 |  |
| hsa04710 | Circadian rhythm - mammal | 3 | PER1 CSNK1D PER2 | | 0 |  | | 0 |  |
| hsa04920 | Adipocytokine signaling pathway | 3 | PPARA STAT3 ADIPOR1 | | 0 |  | | 0 |  |
| hsa04662 | B cell receptor signaling pathway | 1 | PIK3CD | | 0 |  | | 0 |  |
| hsa00330 | Arginine and proline metabolism | 1 | P4HA1 | | 0 |  | | 0 |  |
| hsa04650 | Natural killer cell mediated cytotoxicity | 3 | PAK1 PIK3CD TNFRSF10B | | 0 |  | | 0 |  |
| hsa00052 | Galactose metabolism | 3 | PFKM B4GALT1 PFKL | | 0 |  | | 0 |  |
| hsa04260 | Cardiac muscle contraction | 1 | ATP1B2 | | 1 | ATP1A2 | | 0 |  |
| hsa00520 | Amino sugar and nucleotide sugar metabolism | 2 | PMM1 CYB5R3 | | 0 |  | | 0 |  |
| hsa04350 | TGF-beta signaling pathway | 4 | EP300 BMP7 PPP2CB PITX2 | | 0 |  | | 0 |  |
| hsa04144 | Endocytosis | 11 | TFRC CLTCL1 DNM1 EPN2 PIP5K1A ERBB3 ADRB2 AGAP1 ARAP1 EPN1 AP2B1 | | 0 |  | | 1 | FGFR2 |
| hsa04660 | T cell receptor signaling pathway | 4 | PAK1 PIK3CD PAK4 TEC | | 0 |  | | 0 |  |
| hsa04972 | Pancreatic secretion | 4 | ADCY5 ATP1B2 ATP2A3 ADCY6 | | 1 | ATP1A2 | | 0 |  |
| hsa00030 | Pentose phosphate pathway | 3 | TALDO1 PFKM PFKL | | 0 |  | | 0 |  |
