## Supplementary material for "PPARdelta signaling activation improves metabolic and contractile maturation of human pluripotent stem cell-derived cardiomyocytes": Table2

| **Processes** | | **Immature -> Mature** | **LCFA + GW0742** | **Previously reported** |
| --- | --- | --- | --- | --- |
| **Morphology** | Sarcomere Organization | Improved alignment | Improved, aligned along axis of cell | (Chong et al., 2014; Correia et al., 2017; Feyen et al., 2020; Funakoshi et al., 2021; Giacomelli et al., 2020; Hu et al., 2018; Kamakura et al., 2013; Kosmidis et al., 2015; Leonard et al., 2018; Nunes et al., 2013; Ogasawara et al., 2017; Pasquier et al., 2017; Poon et al., 2015, 2020; Ronaldson-Bouchard et al., 2018; Shadrin et al., 2017; Thavandiran et al., 2013; Tiburcy et al., 2017; Yang et al., 2014, 2019) |
|  | Cell Area | Increased | Increased | (Funakoshi et al., 2021; Kamakura et al., 2013; Nakano et al., 2017; Ogasawara et al., 2017; Ronaldson-Bouchard et al., 2018; Shadrin et al., 2017; Tiburcy et al., 2017; Yang et al., 2014, 2019) |
| **Metabolism** | LCFA Transporters | Increased | Increased CD36 cell surface expression | (Funakoshi et al., 2021; Poon et al., 2020) |
|  | Mitochondrial Content | Increased | Increased | (Feyen et al., 2020; Hu et al., 2018; Lopez et al., 2021; Nakano et al., 2017; Poon et al., 2020; Ronaldson-Bouchard et al., 2018) |
|  | Mitochondrial Morphology | Larger, more peripheral | Larger, more peripheral | (Correia et al., 2017; Funakoshi et al., 2021; Poon et al., 2015, 2020) |
|  | OXPHOS and Active FAO | Increased OXPHOS and Active FAO | Increased OXPHOS and Active FAO | (Correia et al., 2017, 2018; Feyen et al., 2020; Funakoshi et al., 2021; Giacomelli et al., 2020; Hu et al., 2018; Lopez et al., 2021; Nakano et al., 2017; Poon et al., 2020; Ronaldson-Bouchard et al., 2018; Yang et al., 2014, 2019) |
| **DNA Synthesis, Multinucleation and Ploidy** | Proliferation | Reduced | NS | (Funakoshi et al., 2021; Nakano et al., 2017; Poon et al., 2020; Shadrin et al., 2017; Yang et al., 2014) |
|  | Multinucleation | Increased | Increased | (Correia et al., 2017; Funakoshi et al., 2021; Poon et al., 2020) |
|  | Ploidy | Increased | NA |  |
| **Cardiac Gene Expression** | Structural (myofibril and junction proteins) | Increased | NS | (Correia et al., 2017, 2018; Feyen et al., 2020; Funakoshi et al., 2021; Giacomelli et al., 2020; Hu et al., 2018; Kamakura et al., 2013; Leonard et al., 2018; Li et al., 2018; Lopez et al., 2021; Nakano et al., 2017; Poon et al., 2020; Ronaldson-Bouchard et al., 2018; Shadrin et al., 2017; Thavandiran et al., 2013; Tiburcy et al., 2017) |
|  | Channels (Na+, K+, Ca2+ etc) | Increased | NS | (Correia et al., 2017; Feyen et al., 2020; Funakoshi et al., 2021; Hu et al., 2018; Li et al., 2018; Lopez et al., 2021; Nakano et al., 2017; Poon et al., 2020; Shadrin et al., 2017; Tiburcy et al., 2017) |
|  | Metabolic | Increased | Increased expression: - LCFA transporter genes - Metabolic Enzymes | (Correia et al., 2017, 2018; Feyen et al., 2020; Funakoshi et al., 2021; Hu et al., 2018; Lopez et al., 2021; Poon et al., 2015, 2020; Ronaldson-Bouchard et al., 2018; Shadrin et al., 2017; Yang et al., 2019) |
| **Electrophysiology and Contractile Force** | Calcium Handling | Increased Upstroke Velocity, Amplitude or Vmax Decay, etc. | Increased calcium transients duration (20th percentile) | (Correia et al., 2017; Feyen et al., 2020; Funakoshi et al., 2021; Gao et al., 2018; Giacomelli et al., 2020; Hu et al., 2018; Kosmidis et al., 2015; Leonard et al., 2018; Murphy et al., 2021; Nakano et al., 2017; Nunes et al., 2013; Parikh et al., 2017; Poon et al., 2020; Ronaldson-Bouchard et al., 2018; Yang et al., 2014, 2019) |
|  | Action Potentials | Increased amplitude, upstroke velocity, increased action potential duration, etc. | Increased action potential duration (20th and 50th percentiles) | (Correia et al., 2017, 2018; Feyen et al., 2020; Gao et al., 2018; Giacomelli et al., 2020; Kosmidis et al., 2015; Nunes et al., 2013; Poon et al., 2015, 2020; Ronaldson-Bouchard et al., 2018; Shadrin et al., 2017; Tiburcy et al., 2017; Yang et al., 2019) |
|  | Conduction Velocity | Increased | NS | (Nunes et al., 2013; Shadrin et al., 2017) |
|  | Electrical Coupling | Positive force-frequency relationship; increased maximum capture frequency | NS | (Chong et al., 2014; Leonard et al., 2018; Nunes et al., 2013; Pasquier et al., 2017; Ronaldson-Bouchard et al., 2018; Shadrin et al., 2017; Thavandiran et al., 2013; Tiburcy et al., 2017) |
|  | Contractile Force | Increased developed force | Increased systolic and diastolic force; NS developed force | (Feyen et al., 2020; Funakoshi et al., 2021; Gao et al., 2018; Hu et al., 2018; Kosmidis et al., 2015; Leonard et al., 2018; Poon et al., 2015; Ronaldson-Bouchard et al., 2018; Shadrin et al., 2017; Tiburcy et al., 2017; Yang et al., 2014, 2019) |
